# A key interfacial residue identified with in-cell structure characterization of a class A GPCR dimer

**DOI:** 10.1101/664094

**Authors:** Ju Yang, Zhou Gong, Yun-Bi Lu, Chan-Juan Xu, Tao-Feng Wei, Men-Shi Yang, Tian-Wei Zhan, Yu-Hong Yang, Li Lin, Jian-Feng Liu, Chun Tang, Wei-Ping Zhang

## Abstract

G protein coupled receptors (GPCRs) have been shown homo-dimeric. Despite extensive studies, no single residue has been found essential for dimerization. Lacking an efficient method to shift the monomer-dimer equilibrium also makes functional relevance of GPCR dimer elusive. Here, using fluorescence lifetime-based imaging for distance measurements, we characterize the dimeric structure of GPR17, a class A GPCR, in cells. The structure reveals transmembrane helices 5 and 6 the dimer interface, and pinpoints F229 a key residue, mutations of which can render GPR17 monomeric or dimeric. Using the resulting mutants, we show that GPR17 dimer is coupled to both Gα_i_ and Gα_q_ signaling and is internalized, whereas GPR17 monomer is coupled to Gα_i_ signaling only and is not internalized. We further show that residues equivalent to F229 of GPR17 in several other class A GPCRs are also important for dimerization. Our findings thus provide fresh insights into GPCR structure and function.

---

The 800+ G-protein coupled receptors (GPCRs) constitute an important family of membrane receptor. GPCRs are responsible for myriad aspects of cellular functions and make up more than 30% of current drug targets (***Hauser et al., 2018***). A GPCR is characterized with seven transmembrane helices (TM1 to TM7), with its intracellular loops interacting with G proteins and other signaling proteins to transduce signals. Experimental evidences have shown that GPCRs can form dimers and high-order oligomers (***Lohse, 2010***). Indeed, it is widely accepted that class C GPCRs, exemplified by GABA_B_ and metabotropic glutamate receptors, form constitutive dimers. The class C GPCR is characterized with a large extracellular and dimeric N-terminal domain (NTD) for ligand binding (***Muto et al., 2007***). The presence of such a large NTD makes it easy to introduce fluorophores for resonance energy transfer (RET) measurements at the cell surface (***Maurel et al., 2008***). The oligomeric arrangement of reconstituted metabotropic glutamate receptor has also been visualized *in vitro* using cryo-electron microscopy (cryoEM) (***Koehl et al., 2019***).

Different from the class C GPCRs, a class A GPCR contains a much shorter and often unstructured N-terminal segment. It has been known for many years, that β_2_-adrenergic receptor and M1 muscarinic acetylcholine receptor, two class A GPCRs, can dimerize and oligomerize (***Hebert et al., 1996; Hern et al., 2010***). Though increasing evidences have indicated that many class A GPCRs exist in dimers and oligomers, the molecular basis and functional relevance of GPCR dimer/oligomer remains to be fully established (***Lambert and Javitch, 2014; Milligan et al., 2019***). This is because the structures of the GPCR dimers and oligomers are mostly characterized *in vitro* for reconstituted proteins, and also because the transfection and over-expression of an exogenous GPCR in cells may inadvertently promote receptor dimerization and oligomerization. Moreover, a GPCR may dynamically interconvert between monomer, dimer and oligomer, yet an efficient method to shift this equilibrium is lacking (***Kasai and Kusumi, 2014; Gibert et al., 2017***). As such, a structural model of GPCR dimer in the cell membrane would shed light on the structural basis of dimerization and the distinct functions of monomer and dimer.

More than 300 atomic-resolution structures are now available for over 60 unique GPCRs. Despite the technological advance of cryoEM (***Garcia-Nafria et al., 2018***), X-ray crystallography remains the workhorse for structure determination of class A GPCRs. Moreover, some GPCRs have been crystalized as a dimer, either in a single asymmetric unit or related by crystallographic symmetry. Stenkamp analyzed 215 GPCR crystal structures from the protein data base (PDB), and found 13 GPCR dimers with 31 unique dimer interfaces are related by a two-fold rotational *C*_2_ symmetry (***Stenkamp, 2018***), with the interfaces involving TM1/TM2, TM3-TM5, TM3-TM6 or TM5/TM6. However, it is unclear whether these 31 dimer structures represent the physiological quaternary arrangements of the GPCRs. This is because the GPCRs were reconstituted in the detergent, liquid cubic phase or nanodisc for crystallization (***Xiang et al., 2016; Denisov and Sligar, 2017***), and because the majority of the dimer orientations in crystal other than the *C*_2_ symmetry are antiparallel, crisscross and ones incompatible with the cell membrane environment.

Though providing little structural details, a variety of other experimental methods have been employed to assess GPCR dimerization and oligomerization at the cellular level (***Guo et al., 2017; Milligan et al., 2019***). The methods include biochemical approaches like cross-linking, co-immunoprecipitation and protein-fragment complementation (***Romei and Boxer, 2019***). With the fluorophores site-specifically introduced, RET techniques including fluorescence resonance energy transfer (FRET), fluorescence lifetime imaging microscopy FRET (FLIM-FRET), time-resolved FRET (TR-FRET), and bioluminescence resonance energy transfer (BRET) have been frequently used (***Faklaris et al., 2015***). In addition, fluorescence correlation spectroscopy (FCS), spatial intensity distribution analysis (SpIDA) and other single-molecule imaging techniques have revealed membrane co-localization of the receptors (***Calebiro and Sungkaworn, 2018; Gurevich and Gurevich, 2018***). Based on a single FRET distance measurement, it was also possible to model the structure of GPCR dimer in the cell membrane (***Greife et al., 2016***).

With the knowledge about the GPCR dimer, studies were carried out to disrupt GPCR dimerization and to modulate GPCR functions. For example, peptides derived from one of the transmembrane helices have been used to disrupt the dimerization of rhodopsin (***Jastrzebska et al., 2015***), and to inhibit dimerization and activation of β_2_ adrenergic receptor (***Hebert et al., 1996***). GPCR dimerization can also be disrupted with point mutations. For example, Capra et al. introduced single-residue mutations to TM1 of thromboxane receptor and found a small reduction in GPCR dimerization (***Capra et al., 2017***). Based on the protomer structure of M_3_ muscarinic receptor, McMillin et al. systematically mutated all lipid-facing residues in M_3_ muscarinic receptor to alanine, and found only a modest inhibition on GPCR dimerization (***McMillin et al., 2011***). Thus, though GPCR dimerization may involve a small interface (***Baltoumas et al., 2016***), no single residue has been identified essential for dimerization.

GPR17 is a class A GPCR involved in ischemic injuries of kidney, heart, and brain, and also involved in demyelination process associated with multiple sclerosis (MS) and other neurological diseases (***Marucci et al., 2016; Bonfanti et al., 2017; Lu et al., 2018***). GPR17 is phylogenetically related to purinergic P2Y and cysteinyl leukotriene (CysLT) receptors (***Marucci et al., 2016***). Both uracil nucleotides (e.g. UDP-glucose) and cysteinyl leukotrienes (e.g. LTD_4_) can activate GPR17, leading to the inhibition of adenylyl cyclase and the activation of phospholipase C (PLC) (***Ciana et al., 2006; Buccioni et al., 2011***). Moreover, prolonged treatment of UDP-glucose and LTD_4_ can lead to the down-regulation of GPR17 signal, which has been linked to ERK1/2 activation (***Daniele et al., 2014***) and GPR17 receptor internalization (***Daniele et al., 2011***).

To assess whether and how GPR17 homo-dimerizes, we performed FLIM-FRET analysis between GPR17 protomers expressed at the cell membrane. Using a confocal microscopy setup equipped with picosecond pulsed laser, the FLIM-FRET measures the fluorescence lifetime of donor fluorophore, which has an <*r^-6^*> relationship with the distance between donor and acceptor fluorophores (***Becker, 2012; Sun et al., 2013***). Importantly, the fluorescence lifetime is independent of the concentrations of donor and acceptor, and is largely insensitive to the leakage from donor emission and direct excitation of the acceptor. As such, the FLIM-FRET provides a more accurate distance measurement between fluorophores than the standard intensity-based FRET measurement. In addition to N-terminal labeling, we also introduced fluorophores at the extracellular loops (ECLs) of GPR17 using a split-GFP strategy (***Jiang et al., 2016***), and measured multiple FRET distances. Using the distance restraints, we have obtained a well-converged structural model of GPR17 homo-dimer in the cell membrane and identified TM5/TM6 as the dimer interface. Importantly, single point mutations to an essential residue in TM5 can shift the GPR17 monomer-dimer equilibrium, and consequently modulate the downstream signals.

## Results

### GPR17 dimerizes and oligomerizes in the cell membrane

We extracted proteins from mouse tissues, and performed Western blotting analysis for endogenous GPR17 using anti-GPR17 antibody. Under denaturing conditions, a band with the size of GPR17 monomer could be detected (*Figure 1a*). When the samples were prepared without SDS and without boiling, however, GPR17 appears mostly as dimer and high-order oligomers, with only a faint band corresponding to the monomer (*Figure 1b*). Thus, GPR17 from the tissues readily forms dimers and oligomers that can be preserved under non-denaturing conditions.

**Figure 1.**
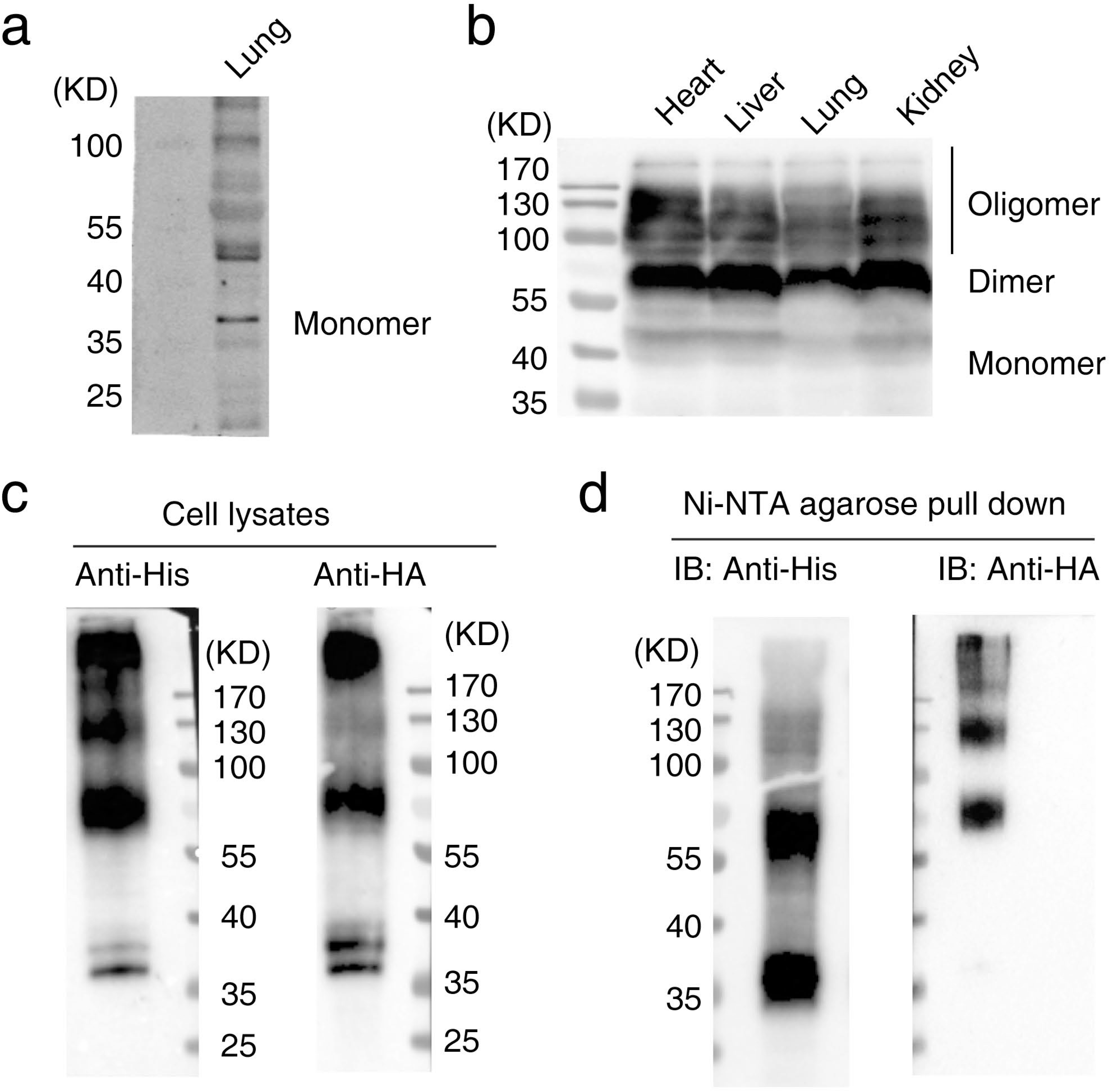
GPR17 dimerizes in mouse tissues and in HEK293 cells. (a) Western blotting analysis of GPR17 from mouse lung under denaturing conditions. 100 μg protein sample each was prepared with 8% SDS sample buffer and was boiled for 5 min before loading. (b) Western blotting analysis of GPR17 from mouse heart, liver, lung and kidney under non-denaturing conditions. 100 µg protein sample each was prepared with SDS-free sample buffer and loaded without boiling. Experiment was repeated for two times. (c) Western blotting analysis of His- and HA-tagged GPR17 proteins in cell lysate under non-denaturing conditions. Samples (100 µg each) were prepared from HEK293 cells co-transfected with His- and HA-tagged GPR17. Experiment was repeated for two times. (d) Western blotting analysis of His- and HA-tagged GPR17 proteins in Ni-NTA agarose affinity-purified sample under non-denaturing condition (50 µg each). The protein samples were purified by using Ni-NTA agarose from His- and HA-tagged GPR17 co-transfected HEK293 cells. Experiment was repeated for two times.

We also expressed human GPR17 proteins in HEK293 cells upon transient transfection. When co-transfected, both His-tagged and HA-tagged GPR17 expressed well, and the monomer, dimer and oligomers of GPR17 could be identified using anti-His or anti-HA antibodies under non-denaturing conditions (*Figure 1c*). Importantly, His-tag and HA-tag can both be identified in GPR17 dimers. Purified from Ni-NTA agarose beads, the His-tagged GPR17 proteins can be blotted with anti-His antibody as monomer and dimers, but can only be blotted with anti-HA antibody as dimers and oligomers (*Figure 1d*). Taken together, the co-immunoprecipitation data indicate that GPR17 homo-dimerizes in the cell membrane.

### GPR17 protomers can FRET in the homo-dimer

To analyze how GRP17 dimerizes, we performed FLIM-FRET measurement between the two protomers of GPR17 at the cell membrane. We engineered GFP and mCherry tags at specific sites of GPR17, as fluorescent donor and acceptor, respectively. The mCherry was appended at the N-terminus of GPR17, and the GFP was either appended at the N-terminus or inserted to one of ECLs using the split-GFP stratagem previously described (***Jiang et al., 2016***). In this labeling approach, the two C-terminal β-strands of GFP (GFP_10-11_) were engineered at ECL1 after residue G128, at ECL2 after residue R214, or at ECL3 after residue R291. Glycine residues are padded at each end of the inserted β-strands, so that the split-GFP can rapidly reorient with respect to GPR17 (*Figure 2—figure supplement 1a, b*). Subsequently, a recombinantly prepared, proteolytically derived GFP fragment comprising the first nine β-strands (GFP_1-9_) was added to the cells, which complements the two β-strands already inserted and regenerates GFP fluorescence. This scheme allowed us to introduce GFP fluorophore to GPR17 only at extracellular surface of the cell membrane. More importantly, a total of four pairs of FRET distances could be obtained using this labeling approach (*Figure 2a*).

**Figure 2.**
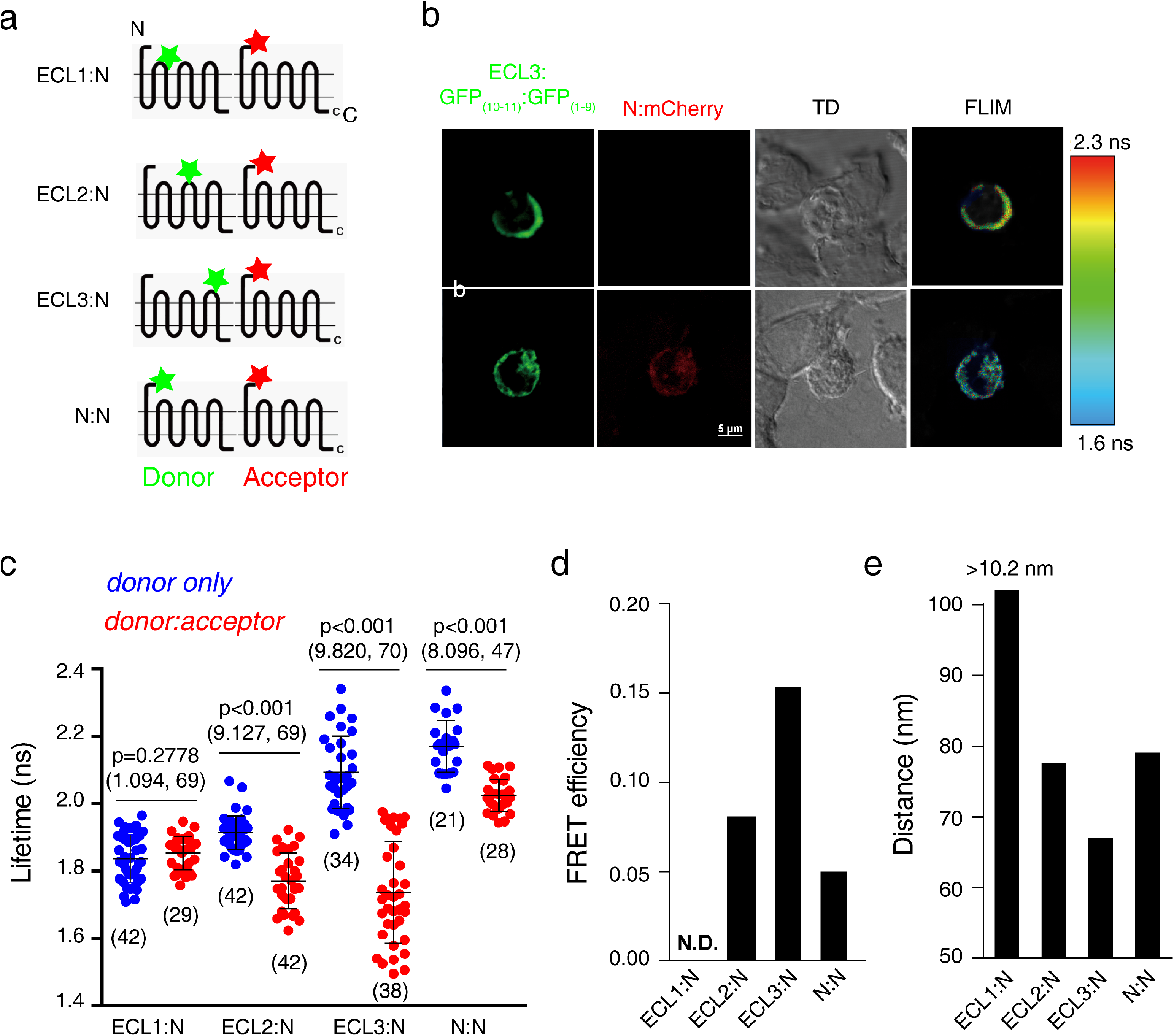
FLIM-FRET measures GPR17 inter-protomer distances in cell membrane. (a) Fluorescence labeling scheme used for FLIM-FRET measurement. Fluorescence acceptor mCherry is fused at the N-terminus, while fluorescence donor GFP is introduced either at the N-terminus or one of the extracellular loops (ECL1-3) using a split GFP stratagem. (b) Representative images showing fluorescent intensity and fluorescent lifetime with GFP (green) inserted at ECL3 of GPR17 with and without mCherry (red) co-expressed. (c) The lifetime of GFP determined from individual HEK293 cells. “*donor only*” means the lifetime of GFP in cells express GFP-tagged GPR17 only. “*donor:acceptor*” means the lifetime of GFP in cells co-express both GFP- and mCherry-tagged GPR17. Mean ± SD. The n value was denoted in the figure under each cluster. Unpaired *t*-test was used for the statistical analysis, and the p value as well as (t, df) were denoted above each cluster. (d) Averaged FRET efficiencies for the four pairs of fluorophores, determined from the decrease of GFP fluorescence lifetime shown in (c). (e) The averaged FRET distances, converted from (d) using a Förster distance of 5.1 nm.

All GFP-tagged GPR17 constructs expressed well and fluoresce. We measured fluorescence lifetime of the GFP alone (τ_D_) at the cell membrane (*Figure 2b*), and found that the averaged lifetime for GFP varies from 2.18 ns to 1.82 ns (*Figure 2c*). The variation is likely a result of the local environment at the insertion site around the fluorophore, which has been noted before (***Suhling et al., 2002; Ito et al., 2009; Berezin and Achilefu, 2010***). The co-transfection of mCherry-tagged GPR17 shortened the fluorescence lifetime of the GFP labeled at another GRP17 protomer. This means that two GPR17 protomers interact with each other, allowing the FRET to occur. Depending on the labeling site, the fluorescence lifetime of GFP with mCherry as FRET acceptor (τ_DA_) varies (*Figure 2c; Table 1*). The largest FRET was observed for GFP introduced at ECL3, whereas essentially no FRET was observed for GFP labeled at the ECL1 of GPR17 (*Figure 2d*). The averaged FRET efficiencies could be converted to distances (*Figure 2e*; *Table 1*). For the ECL1 site, the lack of FRET could be attributed to a large separation between the two fluorophores, and an arbitrary distance (> 10.2 nm, for two times of the Förster distance for GFP-mCherry pair) was given. Since the FRET is related to the inter-fluorophore distance by inverse sixth power and quickly disappears at longer distance, the FLIM-FRET measurements should mostly manifest the arrangement of GPR17 dimer but not the oligomer.

**Table 1.** FRET distances from the measurement of GFP fluorescence lifetime

| FRET pairs | $\tau_D$ (ns) | $\tau_{DA}$ (ns) | p value* | FRET efficiency | FRET distance $r^\dagger$ | Calculated Inter-fluorophore |
| --- | --- | --- | --- | --- | --- | --- |

| | | | | $E^{\P}$ | | distance$^{\ddagger}$ |
| --- | --- | --- | --- | --- | --- | --- |
| ECL1:N | 1.837±0.071 | 1.854±0.049 | 0.28 | 0 | > 102 Å | 104.2 Å |
| ECL2:N | 1.914±0.083 | 1.771±0.083 | 1.75E-13 | 0.075 | 77.5 Å | 77.4 Å |
| ECL3:N | 2.076±0.126 | 1.744±0.157 | 8.44E-15 | 0.16 | 67 Å | 56.1 Å |
| N:N | 2.17±0.077 | 2.025±0.049 | 1.83E-10 | 0.067 | 79 Å | 89.0 Å |

\* Compared between $\tau_D$ and $\tau_{DA}$ , unpaired $t$ -test.
$^{\P}$ Only the average FRET efficiency is given without the S.D. due to large variation for different pixels.
$^{\ddagger}$ Converted from FRET efficiency using the GFP-mCherry Förster distance of 51 Å (*Albertazzi et al., 2009; Akrap et al., 2010*).
$^{\ddagger}$ The averaged distance between the centers-of-mass of the tagged florescent proteins are calculated for the GPR17 homo-dimer structures.

### Modeling the native GPR17 dimer structure pinpoints key interfacial residues

The four pairs of GFP-mCherry tags have different FRET efficiencies and FRET distances (*Table 1*). This can only be explained by that the two GPR17 protomers in the dimer adopt a preferred orientation in the cell membrane. Since there is no experimentally determined structure of GPR17, we first built the model of GPR17 monomer with threaded homology modeling approach (***Roy et al., 2010***) (*Figure 3—figure supplement 1a*). The predicted structures of GPR17 monomer were highly converged (*Figure 3 —figure supplement 1b*). Moreover, the monomer structure remained stable after extended MD simulations in the lipid bilayer environment (*Figure 3—figure supplement 1c-e*), indicative of a correctly folded structure.

**Figure 3.**
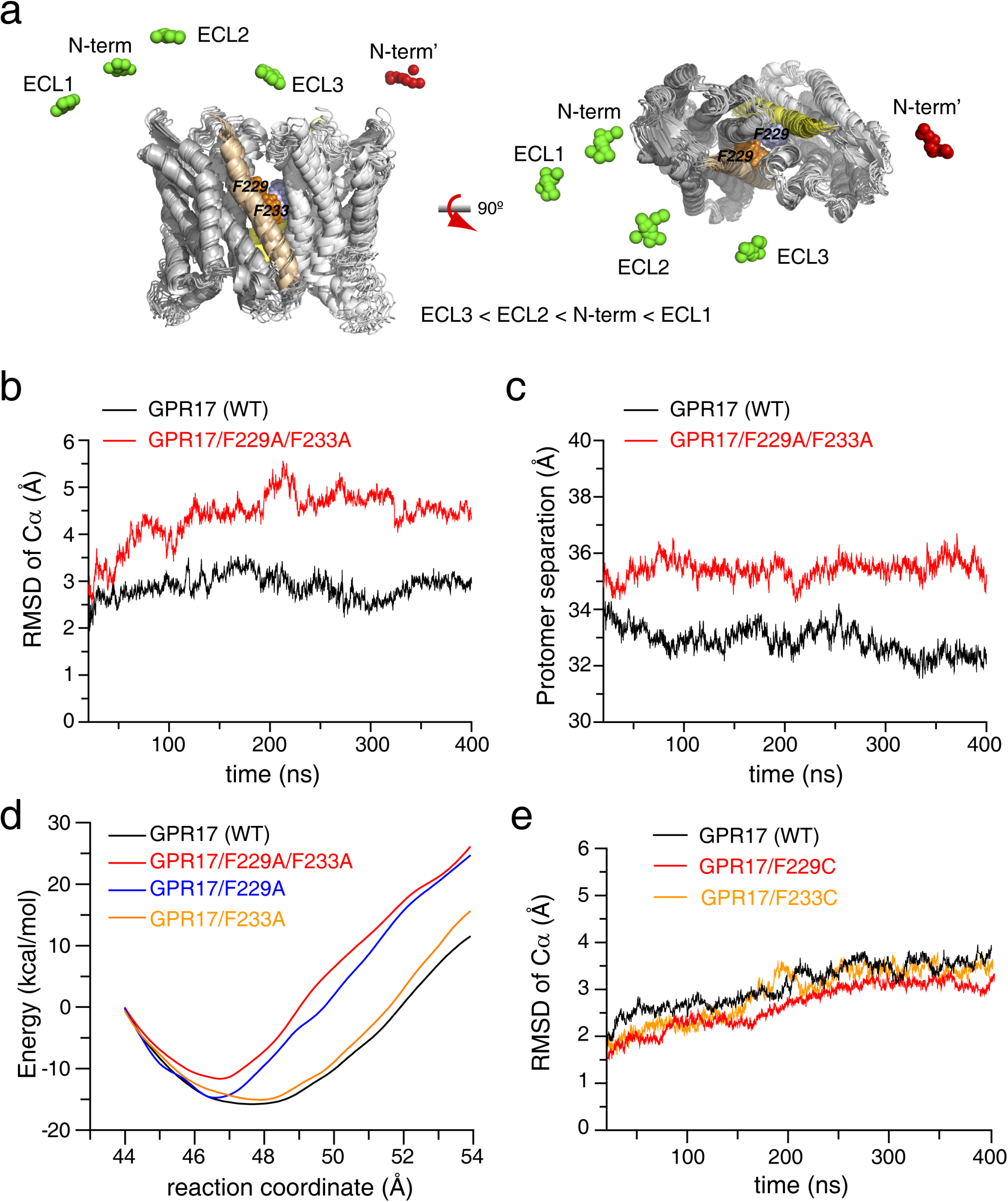
Modeling the structure of GPR17 homo-dimer in cell membrane allows the identification of key interfacial residues. (a) GPR17 dimer structure modeled from inter-protomer FLIM-FRET distances in two orthogonal perspectives. The centers-of-mass of the mCherry and GFP fluorophores are shown as red and green spheres, respectively. The transmembrane helices 5 (TM5) at the dimer interface are colored yellow, and the side chains of F229 and F233 in TM5 are shown as spheres. (b, c) MD simulations of wildtype and F229A/F233A mutant of GPR17 dimer, showing the RMS deviation (RMSD) of the Cα atoms (b), and concurrent separation between the two protomers, i.e. the distance between the centers-of-mass of each GPR17 (c). (d) Steered MD simulations of wildtype and mutant GPR17 dimers. The potential mean force indicates the energetic differences along the reaction coordinate between initial dimeric and final monomeric states. (e) RMS deviations of Cα atoms of wildtype and disulfide-bonded dimeric mutant of GPR17 during MD simulations.

Starting from the monomer structure, we modeled the dimer structure of GPR17 that can simultaneously account for all inter-protomer FRET measurements (*Figure 3 — figure supplement 2*). The sterically allowed conformers of mCherry and GFP tags were first calculated (*Figure 2—figure supplement 1*), and the centers-of-mass of all these conformers with respect to GPR17 protomer were used for the application of FRET distance restraints. We refined the dimeric structure of GPR17 using distance-restrained rigid-body simulated annealing with concurrent enforcement of the *C*_2_ dimer symmetry. The calculated structures of GPR17 dimer were well-converged, with the largest root-mean-square (RMS) deviation of backbone heavy atoms < 3 Å (*Figure 3a*). The dimer interface involves TM5 and TM6, with a total buried surface area 1615 ± 269 Å^2^. In particular, a pair of phenylalanines, residues F229 and F233 in TM5, were found at the dimer interface (*Figure 3a*), and the hydrophobic interactions and possibly aromatic stacking between these residues are likely important for dimerization. The dimer structure of GPR17 was subjected to MD simulations, which remained largely stable in the lipid bilayer (*Figure 3b, c; Figure 3—figure supplement 3*).

To assess the importance of F229 and F233 for GPR17 dimerization, we introduced mutations to these two interfacial residues. With F229 and F233 mutated to alanines *in silico*, the resulting protomer structure remains stable, and the backbone RMS deviation is comparable to that of the wildtype GPR17 (*Figure 3—figure supplement 4a*). In contrast, the dimeric structure of GPR17 becomes unstable, and the overall RMS deviation is larger (*Figure 3b; Figure 3—figure supplement 4b*). This can be attributed to the dissociation of the two GPR17 protomers, as manifested by the increasing distance between the protomers (*Figure 3c*). Steered MD simulations further indicates that F229A/F233A mutant is about 4 kCal/mol less stable than the wildtype protein (*Figure 3d*).

Single point mutations can also change the stability of GPR17 dimer. Steered MD simulations of single alanine mutants of GPR17 indicate that residue F229 contributes more free energy to GPR17 dimerization than F233 does (*Figure 3d*). As a positive control, we performed MD simulations for F229C and F233C mutants of GPR17, one at a time. With an inter-protomer disulfide bond formed, the cysteine mutation allowed the GPR17 to form a stable covalent dimer. Indeed, the RMS deviations of the covalent GPR17 dimers are comparable to that of wildtype GPR17 dimer (*Figure 3e*; *Figure 3—figure supplement 5*). Taken together, intra-membrane residues F229 and to a lesser degree F233 in TM5 can be the key residues for GPR17 dimerization.

### F229 is essential for GPR17 dimerization

To assess the importance of residues F229 and F233 for the dimerization of GPR17 at the cellular level, we introduced F229A and F233A double mutation and repeated the FLIM-FRET measurement. The data showed that upon the mutation, no FRET was observed between fluorophores introduced at ECL3:N sites (*Figure 2a*), and the fluorescence lifetime of GFP remains the same as that of GFP alone (*Figure 4a*). This means that the mutation abrogates GPR17 homo-dimerization, which is consistent with the MD simulations. Western blotting analysis showed that the monomer population significantly increases upon the F229A/F233A mutation (*Figure 4b*, lane 2). However, there remains a large portion of GPR17 dimeric and oligomeric species, which may be heteromers involving interfaces other than TM5.

**Figure 4.**
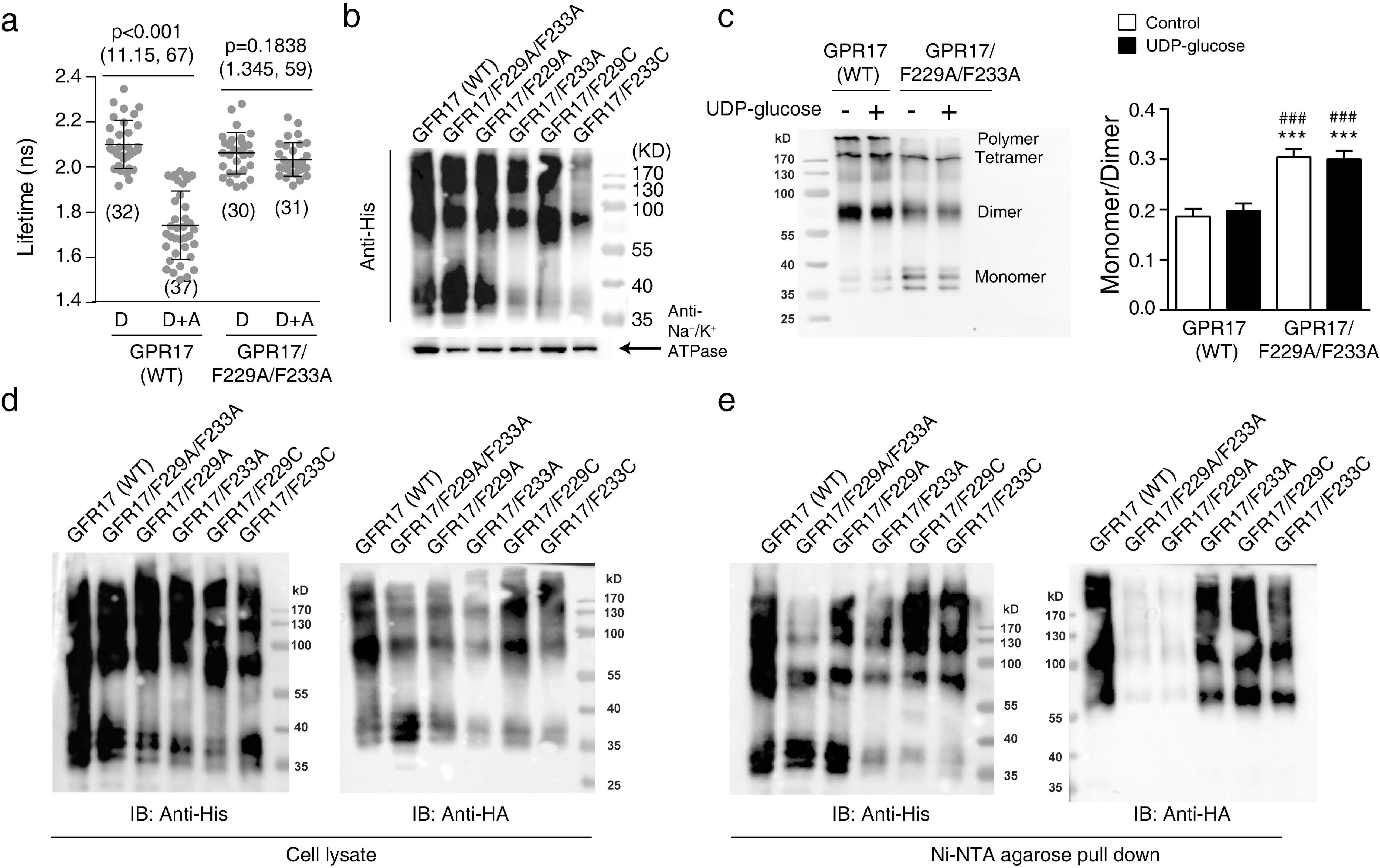
TM5 residue F229 is essential for GPR17 dimerization. (a) The fluorescence lifetime of GFP tagged at ECL3 of GPR17 measured with FLIM-FRET. For F229A/F233A mutant, no difference was observed between “*donor only*” and “*donor:acceptor*”. Mean ± SD. The n value denoted in the figure under each cluster. Unpaired *t*-test was used for the statistical analysis, and the p value and the (t, df) were denoted in the figure above each cluster. (b) Western blotting analysis of His-tagged wildtype and mutant GPR17 proteins expressing in HEK293 cells. 100 mg membrane protein was used for analysis. Experiment was repeated for three times. (c) Effect of UDP glucose on the monomer/dimer ratio between wildtype and F229A/F233A mutant of GPR17. The cells were treated without or with 0.1 mM UDP-glucose for 30 min before harvesting. 100 mg of cell lysate was used for analysis. Mean ± SEM (*Figure 4—source data 1*). n=11. ***p<0.001, compared with GPR17-WT control, the 95% CI of difference was (−0.179 to −0.0566) for the control GPR17/F299A/F233A and (−0.175 to −0.0526) for the UDP-glucose treated GPR17/F299A/F233A. ^###^p<0.001 compared with UDP-glucose treated GPR17-WT, the 95% CI of difference was (−0.168 to −0.0456) for the control GPR17/F299A/F233A and (−0.164 to −0.0416) for the UDP-glucose treated GPR17/F299A/F233A. One-way ANOVA followed by Tukey’s multiple comparisons test was used. p=0.0022 and F(3,28)=6.249 for all group. (d, e) Western blotting analysis using anti-His or anti-HA antibodies for GPR17 proteins extracted from the cell lysate (d) and in Ni-NTA agarose purified sample (e), respectively. His- and HA-tagged wildtype or mutant GPR17 were co-transfected into HEK293 cells. The GPR17(WT) and GPR17/F229A/F233A constructs were repeated for two more times.

To further assess the individual roles of F229 and F233 in GPR17 dimerization, we introduced single point mutations to the protein. Similar to the double mutation, F229A mutation also increased the monomer population of GPR17 (*Figure 4b*, lane 3). The disruption of F233A mutation is smaller, as the F233A mutant of GPR17 remains mainly as dimer and oligomers (*Figure 4b*, lane 4). As expected, F229C and F233C mutations largely eliminate the monomeric species of GPR17, yielding mostly dimer and oligomers (*Figure 4b*, lanes 5 and 6). Interestingly, administration of UDP-glucose, an agonist of GPR17, does not perturb the relative monomer/dimer population of wildtype GPR17 (*Figure 4c*). Similarly, the monomer/dimer ratio for the F229A/F233A mutant is unperturbed upon the addition of UDP-glucose (*Figure 4c*).

To differentiate GPR17 homomers from heteromers, we co-transfected His-tagged and HA-tagged wildtype and mutant GPR17 plasmids. Both anti-His and anti-HA antibodies could blot the monomeric, dimeric and oligomeric species of GPR17 extracted from the cell lysate (*Figure 4d*). On the other hand, anti-HA antibody failed to blot the affinity-purified F229A and F229A/F233A GPR17 mutants (*Figure 4e*, right panel), even though anti-His antibody could blot His-tagged GPR17 proteins purified from Ni-NTA agarose beads (*Figure 4e*, left panel). In comparison, F233A, F229C, and F233C mutants of GPR17 could be blotted with anti-HA antibody as dimers and oligomers, which are similar to the wildtype GPR17. Thus, in agreement with the MD simulations, the experimental data showed that residue F229 in TM5 is essential for the homo-dimerization of GPR17.

### Forced monomer and forced dimer of GPR17 have different functions

GPR17 is coupled to both Gα_i_ and Gα_q_ intracellular signaling pathways (***Ciana et al., 2006; Buccioni et al., 2011***). As mutations to key interfacial residues can make GPR17 monomeric or dimeric, we thus assessed the difference in secondary signals including intracellular cAMP and Ca^2+^ levels. Administration of UDP-glucose to GPR17-expressing cells led to the activation of Gα_i_, which inhibited adenylyl cyclase (AC) activity and inhibited forskalin-induced increase of intracellular cAMP level. Similar to the wildtype GPR17, addition of UDP-glucose also caused significant decrease of the cAMP level for either monomeric or dimeric mutant of GPR17 (*Figure 5a, b*). As such, Gα_i_ signaling is largely unimpaired for the forced monomers and dimers of GPR17.

**Figure 5.**
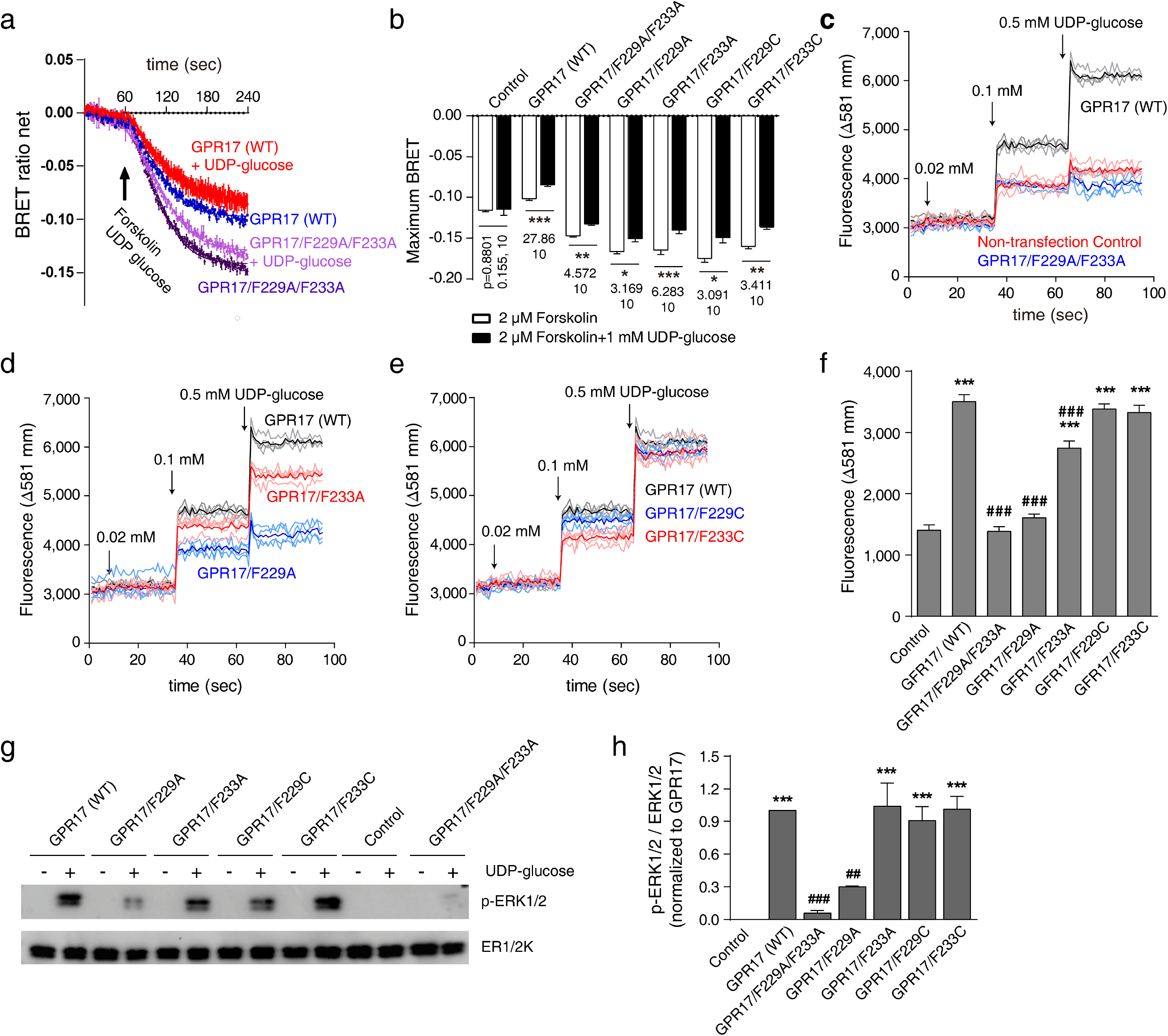
Forced GPR17 monomer and dimer exhibit distinct signaling functions. (a) Representative curves showing BRET ratio of net change, indicative of intracellular cAMP level, upon the administration of forskalin with or without UDP-glucose treatment to GPR17-transfected cells. (b) Statistical analysis of the maximal net change of BRET ratio in (a). mean ± SEM (*Figure 5 — source data 1*). n=6 for each group; \**P* < 0.05, \*\**P*<0.01, \*\*\**P*<0.001, unpaired *t*-test. The value of (t, df) was listed under the histogram bar. (c-e) Fluorescence intensity curves with the sequential administration of 0.02, 0.1 and 0.5 mM UDP-glucose to GPR17-transfected cells, which indicates the intracellular Ca^2+^-levels. For each construct, four curves were recorded for cells from separate wells. The thick curve is the average from four individual curves. The baseline of the curve was adjusted to the similar intensity at 3000. The expression of GPR17 was normalized by the intensity of co-expressed GFP. (f) Statistical analysis of the maximal increase of intracellular Ca^2+^-level induced with the addition of 0.5 mM UDP-glucose. mean ± SEM (*Figure 5 — source data 2*). n=4. \*\*\**P*<0.001, compared with control, the 95% CI of difference was (1660 to 2539) for GPR17(WT), (902.5 to 1781) for GPR17/F233A, (1540 to 2419) for GPR17/229C and (1482 to 2361) for GPR17/F233C; ^###^*P*<0.001, compared with GPR17 (WT) transfected cells, the 95% CI of difference was (1677 to 2555) for GPR17/229A, (1458 to 2336) for GPR17/F233A and (318.9 to 1197) for GPR17/F233A. One-way ANOVA followed by Tukey’s multiple comparisons test was used, with p<0.001 and F(6, 21)=105.3 for all group. (g) Representative images of Western blotting of unphosphorylated and phosphorylated ERK1/2 upon the administration of UDP glucose. (h) Statistical analysis of the ratio of p-ERK over ERK, which indicates the activation of ERK1/2. mean ± SEM (Figure 5*—source data 3*). n=3; \*\*\**P*<0.001, compared with non-transfected control, the 95% CI of difference was (−1.511 to 0.4890) for GPR17(WT), (−0.5692 to 0.4528) for GPR17/F233A, (−0.8096 to 0.2125) for GPR17/229C and (−1.548 to −0.5255) for GPR17/F233C; ^###^*P*<0.01, ^###^*P*<0.001, compared with cells transfected with wildtype GPR17, the 95% CI of difference was (0.4308 to 1.453) for GPR17/229A/F233A and (0.1904 to 1.212) for GPR17/F209A. One-way ANOVA followed by Tukey’s multiple comparisons test was used, with p value <0.001 and F(6, 14)=20.17.

The coupling of Gα_q_ can lead to the activation of phospholipase C (PLC) and subsequent increase of intracellular Ca^2+^ level (***Lu et al., 2018***). We transfected HEK293 cells with a bicistronic vector for simultaneous expression of GPR17 and GFP from a single mRNA transcript. The GFP fluorescence intensities are comparable for the wildtype and mutant GPR17, suggesting similar transfection and expression levels (*Figure 5—figure supplement 1*). Comparing to the non-transfected cells, administration of UDP-glucose to cells transfected with wildtype GPR17 caused an increase of intracellular Ca^2+^ level in a concentration-dependent manner. In contrast, cells transfected with F229A/F233A mutant of GPR17 exhibited almost identical response as the non-transfected cells (*Figure 5c, f*). Similarly, F229A mutation nearly abolished the response of intracellular Ca^2+^ level upon the administration of UDP-glucose. On the other hand, F233A mutation only partially quenched such response (*Figure 5d, f*). For the dimeric control, however, cells transfected with F229C or F233C mutant of GPR17 exhibited almost the same response of intracellular Ca^2+^ level to UDP-glucose treatment as the cells transfected with wildtype GPR17 (*Figure 5e, f*).

Phosphorylation of ERK1/2 was shown downstream of Gα_q_ activation (***Shen et al., 2018***). We found that, administration of UDP-glucose caused phosphorylation of ERK1/2 proteins in GPR17-transfected cells, but not in non-transfected cells (*Figure 5g, h*). For cells transfected with F229A/F233A and F229A mutant of GPR17, the activation of ERK1/2 is significantly lower than the cells transfected with wildtype GPR17. In comparison, the cells transfected with F233A, F229C and F233C mutant of GPR17 exhibited similar levels of ERK1/2 phosphorylation as the cells transfected with wildtype GPR17 (*Figure 5g, h*). Taken together, Gα_q_ signaling involves only the dimeric species of GPR17, which is impaired upon the introduction of monomeric mutations.

### GPR17 is internalized as dimer

It has been shown that GPCRs are internalized as dimer (***Ward et al., 2013; Faklaris et al., 2015***). Therefore, we further assessed the relationship between GPR17 dimerization and receptor internalization. Western blotting analysis showed that, upon the treatment of UDP-glucose, the membrane-associated GPR17 protein level significantly decreased, while at the same time the cytoplasm localized GPR17 significantly increased (*Figure 6a-c*). This means that UDP-glucose can induce GPR17 internalization, even though UDP-glucose does not change the dimer status of GPR17 (*Figure 4c*). In contrast, for the F229A/F33A mutant of GPR17, UDP-glucose does not lead to an increase of cytoplasm-localized GPR17 (*Figure 6a-c*).

**Figure 6.**
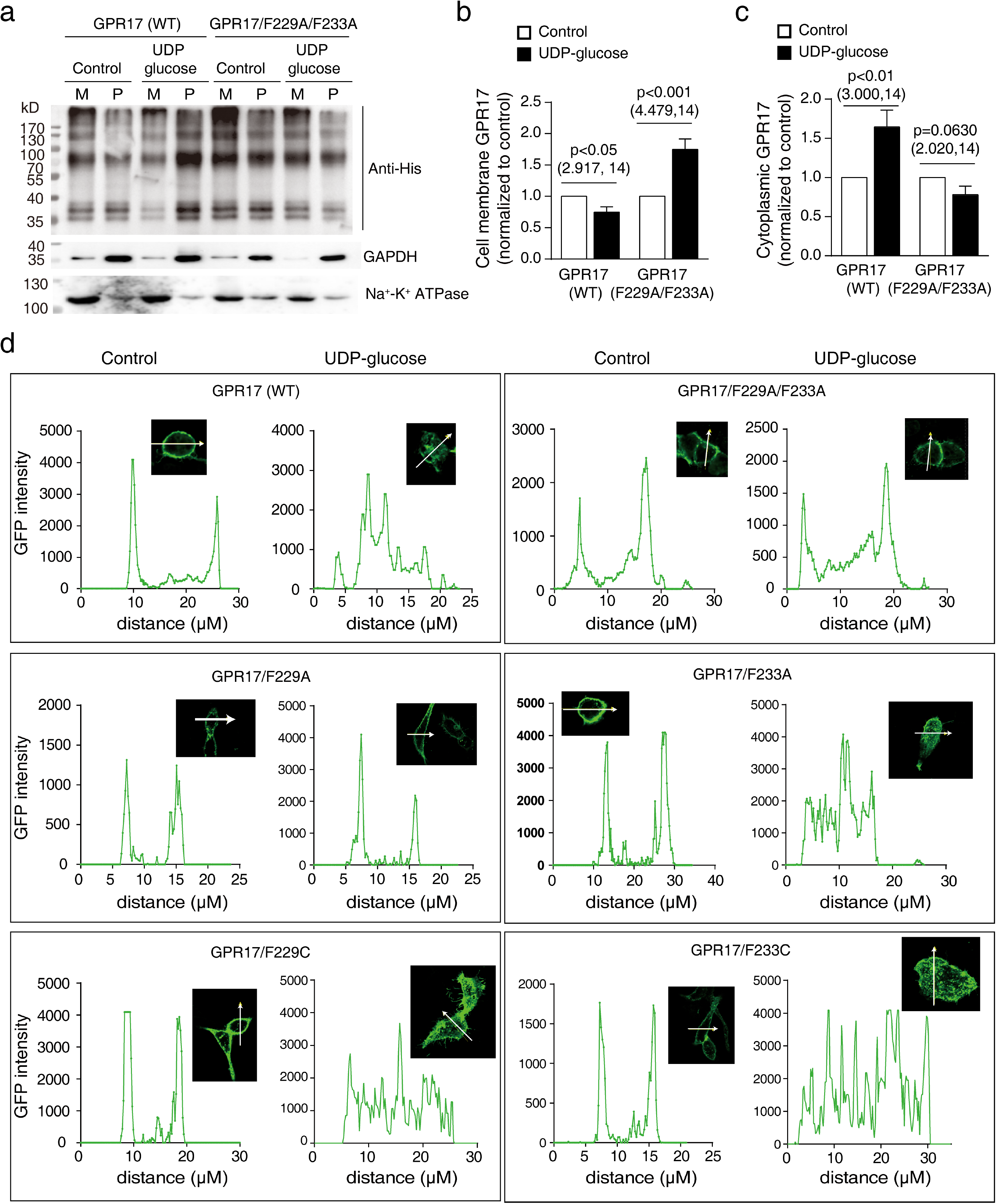
The impact of dimer interface mutations on agonist-induced receptor internalization. (a) Representative Western blot of His-tagged GPR17 and GPR17/F229A/F233A with or without the treatment of 0.1 mM UDP-glucose. Membrane proteins (M) and plasma proteins (P) were prepared separately. (b, c) Statistical analyses of membrane and cytoplasmic proteins for cells treated with or without UDP-glucose for 24 hr. The intensity of cytoplasmic GRP17 is normalized to that of GAPDH, and the intensity of cell membrane GPR17 was normalized to that of Na^+^-K^+^ ATPase. Mean ± SEM (*Figure 6—source data 1*). n=8; unpaired *t*-test. The p value and (t, df) was listed in the figure above the bar. (d) Sectional analysis of fluorescence intensity for cells transfected with GPR17 and labeled with split-GFP. GFP fluorescence was initially observed at the membrane without UDP-glucose, and depending on the GPR17 construct, could be visualized inside the cells following the treatment of UDP-glucose. Experiment was repeated for three times.

To visualize the internalization process, we introduced a split-GFP fluorophore at ECL3 of GPR17 (*Figure 2a*). Thus, only the GPR17 protein localized at the cell membrane can be initially visualized. Note that introduction of the fluorophore using this approach does not affect the trafficking of GPR17 to the cell membrane (*Figure 6d; Figure 6—figure supplement 1*). 24 hours after the treatment of UDP-glucose, most GFP-tagged GPR17 becomes internalized for cells transfected with wildtype, F233A, F229C and F233C mutants of GPR17. In contrast, the F229A/F233A and F229A mutants of GPR17 remain mostly at the membrane (*Figure 6d; Figure 6—figure supplement 1*), which is consistent the Western blot analysis (*Figure 6a-c*). As such, UDP-glucose induced the internalization of GPR17 dimer but not the monomer.

### Residues equivalent to F229 in other GPCRs are also important for dimerization

The µ-opioid receptor has been crystalized as a dimer in each asymmetric unit (***Manglik et al., 2012***), and the dimer also utilizes TM5 and TM6 as the interface (***Baltoumas et al., 2016; Stenkamp, 2018***), involving residue F239 (*Figure 7—figure supplement 1a*). Indeed, the dimer structure of μ-opioid receptor is stable during MD simulations, while an alanine mutation to F239 in helix TM5 drastically decreases the stability of μ-opioid receptor dimer (*Figure 7—figure supplement 1b, c*). Western blotting analysis confirmed that the receptor can form a homo-dimer in cells (*Figure 7a, b*). Moreover, we found that the F239A mutation for the µ-opioid receptor increased the population of the monomeric species, while decreased the population of homo-dimeric and homo-oligomeric species carrying both His-and HA-tags (*Figure 7a, b*).

**Figure 7.**
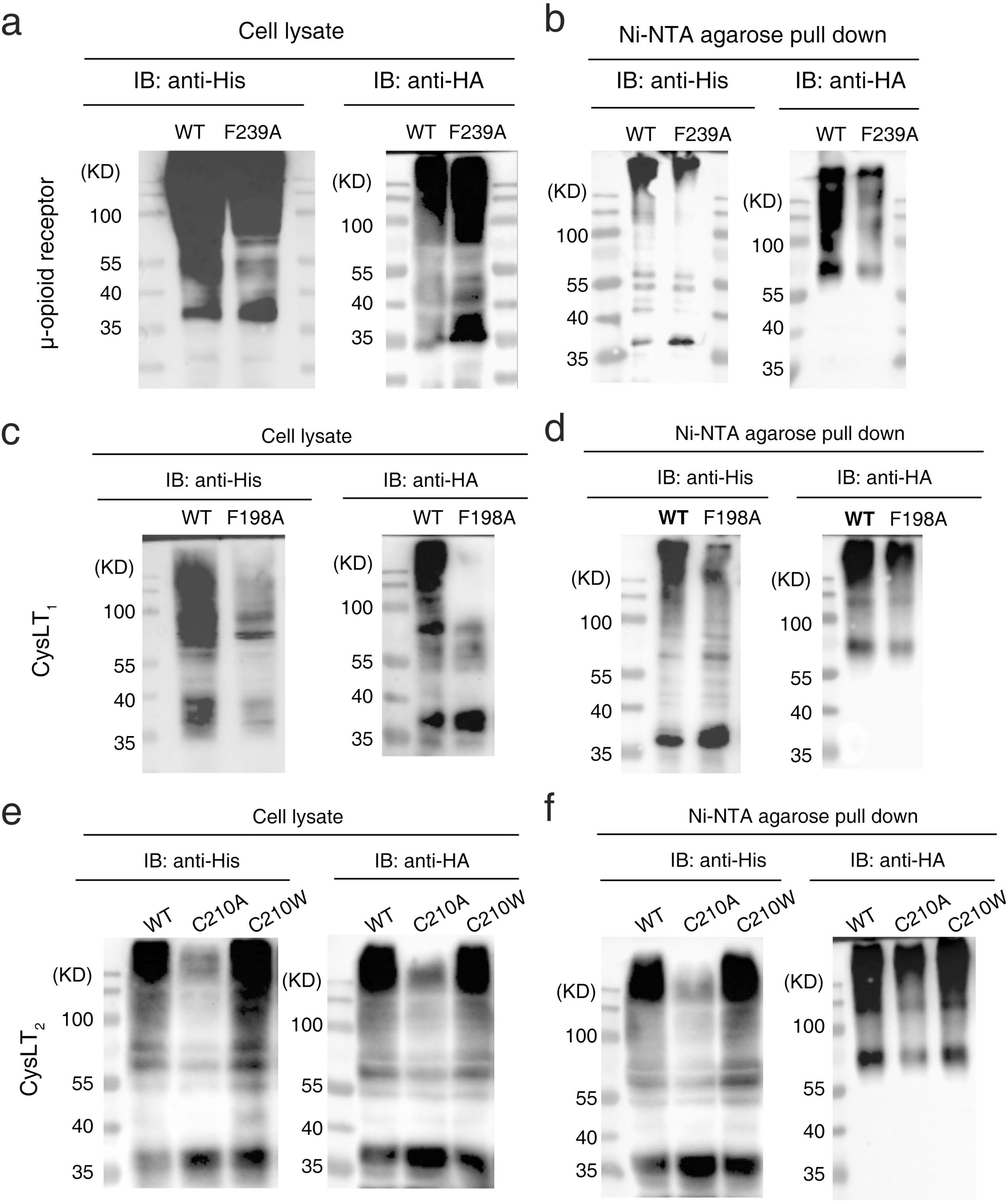
Residues equivalent to F229 in GPR17 is important for the dimerization of several other GPCRs. (a, b) Western blot analysis of µ receptor in cell lysate (a) and in Ni-NTA agarose purified sample (b), respectively. (c, d) Western blot analysis of CysLT_1_ in cell lysate (c) and in Ni-NTA agarose purified sample (d), respectively. (e, f) Western blot analysis of CysLT_2_ in cell lysate and in Ni-NTA agarose purified sample, respectively. 100 μg samples of cell lysate and 50 μg of purified samples were used for Western blotting analysis under non-denature condition. All experiments were repeated for two times.

GPR17 is phylogenetically related to cysteinyl leukotriene receptors including CysLT_1_ and CysLT_2_. Residues F198 in CysLT_1_ and C210 in CysLT_2_ are equivalent to residue F229 in GPR17 (***Isberg et al., 2015; Pandy-Szekeres et al., 2018***). We thus assessed their roles for the dimerization of CysLT_1_ and CysLT_2_ receptors. Co-transfection of His-tagged and HA-tagged CysLT_1_ and CysLT_2_ receptors showed that both receptors can form dimers and oligomers (*Figure 7c, e*). Purified using Ni-NTA agarose beads, the His-tagged proteins can be blotted with anti-His antibody as monomer, dimers, and oligomers, but can only be blotted with anti-HA antibody as dimers and oligomers (*Figure 7d, f*). This means that both CysLT_1_ and CysLT_2_ receptors can form homo-dimers. Importantly, the F198A mutation increased the monomer population of CysLT_1_ (*Figure 7c, d*). Similarly, the C210A mutation but not the C210W mutation increased the population of CysLT_2_ monomer (*Figure 7e, f*). Note that C210W is a naturally occurring mutation for human CysLT_2_ with no debilitating phenotype (***Pandy-Szekeres et al., 2018***), and therefore, it is likely that inter-protomer aromatic stacking between the tryptophan residues promotes CysLT_2_ dimerization in lieu of a covalent disulfide linkage. Interestingly, the F198A mutation in CysLT_1_ and the C210A mutation in CysLT_2_ were found to increase the monomer population at the expense of both dimers and oligomers, whereas the population of homo-dimers and homo-oligomers carrying both His- and HA-tags were found about the same for mutant and wildtype proteins. Thus, residues F198 of CysLT_1_ and C210 of CysLT_2_ are important for receptor homo-dimerization, but they can also be involved in receptor hetero-dimerization.

## Discussion

On the basis of FLIM-FRET analysis, we have modeled the dimeric structure of GPR17, a class A GPCR, in its native cell membrane environment. In doing so, we were able to pinpoint F229 as the key interfacial residue for GPR17 dimerization. Mutating this key residue can force GPR17 monomeric or dimeric, which then allowed us to dissect the respective functions of GPR17 monomer and dimer (*Figure 8*). In addition, residue equivalent to F229 in certain other GPCRs can also be important for receptor dimerization.

**Figure 8.**
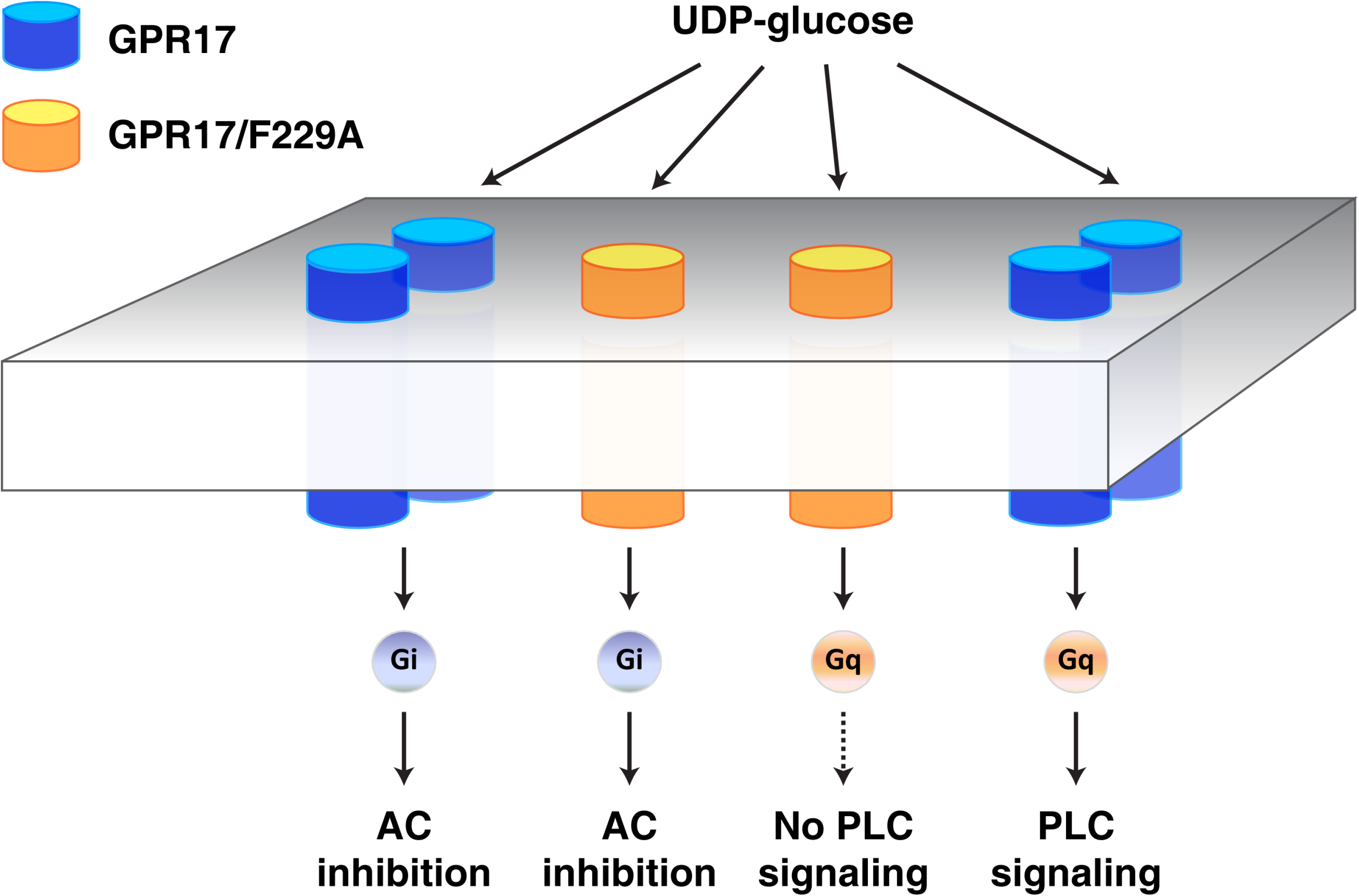
Schematic illustration of the signals of the G-proteins coupled with wildtype and F229A mutant of GPR17 receptors.

Using a split GFP labeling strategy (***Jiang et al., 2016***), we achieved multi-site labeling and obtained several cross-validating distance restraints. The multiple inter-protomer distances, in conjunction with rigid-body simulated annealing refinement, allowed us to obtain a precise description of the GPR17 dimer structure in cells. In comparison, in a previous study with FRET-based modeling of GPCR dimer structure in the cell membrane (***Greife et al., 2016***), only a single distance was obtained between the fluorescent proteins fused at protein N-termini. As a result, several dimeric models involving different interfaces could all account for this FRET distance measurement. Here, the structure of GPR17 dimer is well converged, showing unambiguously that GPR17 dimer interface involves TM5 and TM6.

The well-converged GPR17 dimer structure allowed us to identify a key interfacial residue, F229 in TM5. The importance of this key residue was assessed with a range of experiments and MD simulations. Alanine mutation of F229 makes GPR17 monomeric, whereas cysteine mutation yields an intermolecular disulfide bond and makes GPR17 dimeric. To the best of our knowledge, this is the first time it has been shown, that a single point mutation can effectively shift the equilibrium and force a class A GPCR monomeric or dimeric. Interestingly, the equivalent of F229 is highly conserved among P2Y and CysLT GPCRs. The only two exceptions are human P2Y_11_ and CysLT_2_ receptors, both of which have cysteine residues instead. We have also experimentally confirmed the importance of F198 in CysLT_1_ and C210 in CysLT_2_, the two equivalent residues. We also found that TM5 residue F239 in μ-opioid receptor, a more distantly related class A GPCR, is also important for receptor dimerization.

It has been noted that GPCR dimerization is dynamic. First, the association between GPCR protomers is dynamic, and the GPCR can alternate between monomer, dimer and oligomer. This has been shown for class A receptors including M_1_ muscarinic receptor (***Hern et al., 2010***), and β_1_, β_2_ and GABA_B_ receptors in the cell membrane (***Calebiro et al., 2013***). Secondly, the arrangement of the GPCR dimer is dynamic. It has been shown that the dimer of metatropic glutamate receptor 2, a class C GPCR receptor, utilizes a different interface upon the addition of the agonist (***Xue et al., 2015***). In the present study, the dimer of GPR17 is rather stable, and the addition of the agonist did not change the monomer/dimer ratio. Moreover, F229 is located outside the heptahelical bundle of GPR17, and mutations to F229 have little impact on the structure of GPR17 protomer. Accordingly, the application of UDP-glucose led to the same degrees of inhibition of the intracellular cAMP levels for cells transfected with wildtype GPR17 and with GPR17 mutants. Importantly, the results indicate that GPR17-coupled Gα_i_ signaling and adenylyl cyclase activity are independent of monomer-dimer equilibrium of the receptor.

On the other hand, mutations to residue F229 in TM5 has a large effect on Gα_q_ signaling. Upon the activation of GPR17-coupled Gα_q_, PLC and PKC activities are up-regulated, which then lead to changes in intracellular Ca^2+^-level and ERK1/2 phosphorylation level (***Marucci et al., 2016; Bonfanti et al., 2017; Lu et al., 2018***). Comparing with the cells transfected with wildtype GPR17, cells transfected with GPR17 monomeric mutant exhibits significantly lower intracellular Ca^2+^ and ERK1/2 phosphorylation levels upon UDP-glucose treatment. Thus, GPR17 dimer exhibits a biased signaling from GPR17 monomer. It has been previously reported that GPCR can be internalized as homo-dimer or hetero-dimers (***Ecke et al., 2008; Ward et al., 2013; Ge et al., 2017; Smith et al., 2017***). Consistent with these reports, we found that the monomeric mutations impeded the internalization of GPR17 that could be otherwise triggered upon agonist binding. It can be thus concluded that GPR17 is internalized as a dimer.

Besides residue F229 in TM5, the structural model of GPR17 dimer also indicates that TM5 residue F233 from the two protomers are also outward facing and close to each other. Though F233C mutation could lock GPR17 in a covalent dimer, F233A mutation failed to disrupt the GPR17 dimer. On the contrary, the mutation appears to decrease the monomer population (*Figure 4d, e*), which can be attributed to different and possibly stronger GPR17-GPR17 interactions upon the mutation. Nevertheless, cells transfected with F233A mutant of GPR17 has lower Ca^2+^-level in response to UDP-glucose treatment than the cells transfected with wildtype protein (*Figure 5d, f*). This means that F233 plays a subtler role in receptor dimerization than F229, and probably helps to correctly position F229 side chains and the two GPR17 protomers for dimerization.

In summary, we have identified a key residue in GPR17 essential for receptor dimerization, and dissected the respective functions of GPR17 monomer and dimer. Our data also suggests that GPR17 homo- and hetero-dimerization utilize different interfaces, which can be further investigated. Moreover, the equivalent residues in other class A GPCRs can also be mutated to evaluate to the respective functions of forced monomer and forced dimer. The dissection of GPCR functions by monomer and dimer can also help the development of “smart” drugs for precise pharmacological intervention of GPCR signaling.

## Acknowledgements

Funding: The work has been supported by the National Key R&D Program of China (2018YFA0507700 to Z.G., C.T., Y.B.L., and W.P.Z., and 2016YFA0501200 to Z.G. and C.T.), and by the National Natural Science Foundation of China (81573400 to W.P.Z., 91753132 and 31770799 to C.T., and 31400735 to Z.G.).

## Ethics

Ethics Animal experimentation: The mouse was handled in accordance with the Guideline for the Care and Use of the Laboratory Animals of the National Institutes of Health. All procedures were approved by the Ethics Committee of Laboratory Animal Care and Welfare, Zhejiang University School of Medicine. The experiment was performed strictly according the approved protocols (ZJU2015-12-02).

## Materials and methods

### Construction of plasmids

All plasmids used in this study are listed in Supplementary table 1, and the primers used for the construction of plasmids are listed in Supplementary table 2.

| Reagent type (species) or resource | Designation | Source or reference | Identifiers | Additional information |
| --- | --- | --- | --- | --- |
| Cell line (human) | HEK 293 | Cell Biology of the Chinese Academy of Sciences, Shanghai, China | cat#GNHu43 |  |
| plasmid | pIRES2-EGFP | Clontech Laboratories, Mountain View, USA | cat#6029-1 |  |
| plasmid | pcDNA3.1(+) | Invitrogen by Thermo Fisher Scientific, California, USA | cat#V79020 |  |
| GPR17 ligands | UDP-glucose | Sigma-Aldrich Co., St. Louis MO, USA | cat#U4625-400MG |  |
|  | BCA Protein Assay Kit | Beyotime Biotechnology, Shanghai, China | cat#P0009 |  |
| Inhibitor | protease inhibitor cocktail | Beyotime Biotechnology, Shanghai, China | cat#P1008 |  |
| Inhibitor | Phosphatase inhibitor cocktail | Beyotime Biotechnology, Shanghai, China | cat#P1081 |  |
| protein extraction reagent | RIPA Lysis Buffer | Beyotime Biotechnology, Shanghai, China | cat#P0013B |  |
| protein extraction kit | Total Protein Extraction Kit for Cells and Tissues | Kangchen Bio-tech Inc., Shanghai, China | cat#KC-415 |  |
| Protein Extraction Kit | Membrane and Cytosol Protein Extraction Kit | Beyotime Biotechnology, Shanghai, China | cat#P0033 |  |
| Antibody | anti-phosphor-ERK1/2 (mouse polyclonal) | Cell Signaling Technology, Boston, USA | cat#9106 | 1:1000 |
| Antibody | anti-total ERK1/2 (rabbit polyclonal) | Cell Signaling Technology, Boston, USA | cat#9102 | 1:1000 |
| Antibody | anti-His Tag (mouse polyclonal) | Ruiying Biological Technology, Suzhou, China | cat#RLM002 | 1:2000 |
| Antibody | anti-HA Tag (rabbit polyclonal) | Proteintech Group, Wuhan, China | cat#51064-2-AP | 1:500 |
| Antibody | anti-GAPDH (mouse polyclonal) | Proteintech Group, Wuhan, China | cat#60004-1-Ig | 1:4000 |
| Antibody | anti-GPR17 (mouse polyclonal) | Santa Cruz Biotechnology, Texas, USA | cat#SC-514723 | 1:500 |
| Antibody | anti-Na<sup>+</sup>/K<sup>+</sup> ATPase (rabbit polyclonal) | Proteintech Group, Wuhan, China | cat#14418-1-AP | 1:500 |
| Antibody | anti-mouse IgG, HRP-Linked secondary (horse polyclonal) | Cell Signaling Technology, Massachusetts, USA | cat#7076 | 1:3000 |
| Antibody | anti-rabbit IgG, HRP-Linked secondary (goat polyclonal) | Cell Signaling Technology, Massachusetts, USA | cat#7074 | 1:3000 |
| Antifade mountant | anti-fade medium with DAPI | Invitrogen, Carlsbad, CA, USA | cat#P36931 |  |
| reagent |  |  |  |  |
| cAMP assay reagent | Coelenterazine h | Promega, Madison, USA | cat#S2011 |  |
| cAMP assay reagent | Forskolin | Sigma-Aldrich, St. Louis, MO, USA | cat#F6886-10MG |  |
| Calcium assay reagent | Rhod 2 AM | Life technologies, Eugene, USA | cat#R1245MP |  |
| Calcium assay plate | microplate reader | Perkin-Elmer, Waltham, USA | cat#6005290 |  |
| CAMP assay 96 MicroWell | 96-well white plate | Thermo Scientific Nunc, Denmark | cat#236105 |  |
| Calcium assay reagent | Pluronic™ F-127 (20% Solution in DMSO) | Sigma-Aldrich, St. Louis, MO, USA | cat#P3000MP |  |
| Protein purification reagent | Ni-NTA agarose beads | Qiagen, Hilden, Germany | cat#1018244 |  |
| Cell culture medium | DMEM | Gibco by Thermo Fisher Scientific | cat#11995500BT |  |
| Cell culture medium | Opti-MEM | Gibco by Life Technology, Carlsbad, USA | cat#31985-070 |  |
| Cell culture reagent | fetal bovine serum | Zhejiang Tianhang Biotechnology, Hangzhou, China | cat#13011-8611 |  |
| Transfection reagent | Polyethyleneimine, linear, M.W. 25,000 | Alfa Aesar Chemical, Shanghai, China, CAS number: 9002-98-6 | cat#43896 |  |
| Western reagent | Western ECL substrate | Biorad, Hercules, USA | cat#170-5061 |  |
| Western reagent | Immobilon-NC Membrane, 0.45 µm | Millipore Corporation, Billerica, USA | cat#HATF00010 |  |
| Western Imaging System | ChemiDoc™ MP V3 | Biorad, Hercules, USA | cat#17001395 |  |
| Microscope | A1 confocal laser-scanning microscope | Nikon, Tokyo, Japan |  |  |
| Photon counting equipment | Time correlated single photon counting (TCSPC) module | PicoQuant, Berlin, Germany |  |  |
| Diode laser | diode laser | LDH-P-C-485B, PicoQuant, Germany |  |  |
| cAMP assay system | Mithras LB 940 | Berthold, Bad Wildbad, Germany |  |  |
| Software | Quantity One | Biorad, Hercules, USA |  |  |
| Software | SymphoTime 64 | PicoQuant, Germany |  |  |
| Software | GraphPad Prism 6.0 | GraphPad Software |  |  |
| Software | PyMOL Molecular Graphics System Version 2.2 | Schrödinger, LLC. |  |  |

### Cell lines and cell culture

HEK293 cells were purchased from the Institute of Cell Biology of the Chinese Academy of Sciences (Cell Biology of the Chinese Academy of Sciences, Shanghai, China). Cells were grown in Dulbecco’s modified essential medium (DMEM, Gibco by Thermo Fisher Scientific) supplemented with 10% heat-inactivated fetal bovine serum (Zhejiang Tianhang Biotechnology, China) at 37 °C, in the atmosphere of 5% CO_2_.

### Cell transfection

A cost-efficient transfection approach was used with polyethyleneimine (PEI, linear with average molecular weight 25,000 Da, Alfa Aesar Chemical, Shanghai, China) according to the literature (***Longo et al., 2013***). Briefly, PEI stock solution (1 μg/μL) was prepared by dissolving in endotoxin-free ddH_2_O. A suitable number of cells were seeded one day prior to transfection, and as a result the cells were ∼70-80% confluent at the time of transfection. Before transfection, plasmid DNA and PEI (at the weight ratio of 1:2) was carefully mixed with appropriate volume of Opti-MEM (Gibco by Life Technology, Carlsbad, USA), and the mixture was incubated at room temperature for 25 min. The freshly formed DNA/PEI precipitates were carefully pipetted to the cells. The medium containing transfection reagents was removed and fresh medium was added 6 h after transfection.

### Ni-NTA agarose affinity purification

His-tagged and HA-tagged wildtype or mutant receptors (at DNA ratio of 1:1) were transfected to HEK293 cells. 36 hours after transfection, the cells were washed with PBS buffer, lysed for 60 min at 4 °C in lysis buffer (containing 25 mM Tris–HCl pH 8.0, 150 mM NaCl, 1% n-Dodecyl-β-D-maltopyranoside, protease inhibitor cocktail, Beyotime Biotechnology, Shanghai, China), and centrifuged at 14,000 g for 15 min at 4 °C. The supernatant was incubated with Ni-NTA agarose beads (Qiagen, Germany, catalog number 1018244) for 30 min at 4 °C. The Ni-NTA agarose beads were washed three times with the washing buffer (25 mM Tris–HCl pH 8.0, 150 mM NaCl, 0.05% n-Dodecyl-β-D-maltopyranoside, 20 mM imidazole). The His-tagged protein was eluted with the elution buffer (25 mM Tris–HCl pH 8.0, 150 mM NaCl, 0.05% n-Dodecyl-β-D-maltopyranoside, 300 mM imidazole). The protein concentrations of total cell lysate and of Ni-NTA agarose affinity-purified protein were determined using a BCA Protein Assay Kit (Beyotime Biotechnology, Shanghai, China, catalog number P0009).

### Western blotting analysis

One C57BL/6J male mouse was purchased from Zhejiang Academy of Medical Sciences. The mouse brain, lung, heart and kidney tissues were separated after deeply euthanized by intraperitoneal injection of pentobarbital sodium (250 mg/kg). The total protein from the tissues was extracted by using protein extraction kit (Kangchen Bio-tech Inc., Shanghai, China, catalog number KC-415). The total cell lysate was prepared with RIPA buffer (Beyotime Biotechnology, Shanghai, China, catalog number P0013B). The membrane protein and cytoplasmic protein were isolated by Membrane and Cytosol Protein Extraction Kit (Beyotime Biotechnology, Shanghai, China, catalog number P0033). The protein concentration was determined with the use of BCA Protein Assay Kit (Beyotime Biotechnology, Shanghai, China).

Protein samples were separated on a 10% SDS-PAGE and transferred to nitrocellulose membranes (Millipore Corporation, Billerica, USA). The non-denaturing condition was achieved with the use of SDS-free sample buffer and without sample boiling before loading sample. The membranes were blocked with 5% non-fat dried milk at room temperature for 60 min, and were incubated with first antibodies overnight at 4 °C. To detect the expression of native and transfected GPR17 proteins, the first antibodies include mouse anti-GPR17 polyclonal antibody (Santa Cruz Biotechnology, Texas, USA, SC-514723, diluted at 1:500), mouse anti-His tag polyclonal antibody (Ruiying Biological Technology, Suzhou, China, RLM002, 1:2000), anti-HA antibody (Proteintech Group, Wuhan, China, 51064-2-AP, 1:500), mouse anti-GAPDH polyclonal antibody (Proteintech Group, 60004-1-Ig, 1:4000) and rabbit anti-Na^+^/K^+^ ATPase antibody (Proteintech Group, Wuhan, China, 14418-1-AP, 1:500). The membranes were then washed and incubated with the corresponding secondary antibodies for 1.5 hours at room temperature, which include HRP-conjugated anti-mouse IgG (Cell Signaling Technology, Massachusetts, China, 7076, 1:3000) and HRP-conjugated anti-rabbit IgG (Cell Signaling Technology, 7074, 1:3000). After washing with TBS-T buffer (25mM Tris, 137 mM NaCl, 0.1% Tween-20, pH 7.6), the membranes were incubated in ECL substrate (Biorad, Hercules, USA), and the chemiluminescence was detected on ChemiDocTM MPV3 (Biorad, Hercules, USA). Densitometric evaluation of the blots was performed using the program Quantity One (Biorad, Hercules, USA).

To assess the internalization of GPR17, HEK 293 cells were transfected with His-tagged GPR17. Two days after transfection, 100 μM UDP-glucose (Sigma-Aldrich Co., St. louis MO, USA, U4625-400MG) was applied to the culture medium. 24 hours after treatment, the cells were collected, and the cell membrane protein and cytoplasmic protein was separated prior for Western blotting analysis. The density of His-tagged GPR17 in cell cytoplasmic fraction was normalized to that of GAPDH, and the density of GPR17 in membrane fraction was normalized that of Na^+^-K^+^ ATPase. Both the membrane expressing and the cytoplasmic expressing proteins were normalized to the relative control (without UDP-glucose treatment) blotted on the same membrane.

### FLIM-FRET measurement

To perform fluorescence lifetime imaging microscopy, we used GFP as FRET donor and mCherry as FRET acceptor. The GFP was either fused to the N-terminal of GPR17 or labeled at one of the three extracellular loops (ECLs) using split GFP strategy as described previously (***Jiang et al., 2016***). For the internal labeling of GPR17 using the split-GFP stratagem, the last two β-strands of GFP are site-specifically inserted, with four glycine residues flanking the inserted strands for flexibility of the fluorophore. The four pairs of fluorescent donor-acceptor pairs are listed in Table 1.

HEK 293 cells were transfected with donor plasmid alone (for donor only control) or co-transfected with both donor and acceptor plasmids (at DNA ratio of 1:1). 6 hours after transfection, cells were passed into 24-well plates with glass coverslips at the bottom, and were cultured for another 24 hours. For split GFP labeling, the cells were incubated with 2 µM GFP_(1-9)_, a fragment comprising the first 9 β-strands of GFP, in PBS buffer at 37 °C for 20 min. Excess GFP _(1-9)_ fragment was removed by washing twice with PBS. The cells were then fixed with freshly prepared 4% paraformaldehyde at 37 °C for 10 min. The cover slides with fixed cells were mounted on the glass slides with an anti-fade medium containing DAPI (Invitrogen, Carlsbad, CA, USA). The fluorescence intensities of donor and acceptor were determined with an A1 confocal laser-scanning microscope (Nikon, Tokyo, Japan). Images were obtained by sequential excitation at 488 nm for GFP, 561 nm for mCherry and 465nm for DAPI, respectively.

Fluorescence lifetime of donor fluorophore without or with the acceptor fluorophore present was evaluated with an A1 confocal laser-scanning microscope equipped with a time correlated single photon counting (TCSPC) module (PicoQuant, Berlin, Germany). The donor was excited at 485 nm using a picosecond pulsed with a diode laser (LDH-P-C-485B, PicoQuant, Germany) at 40 MHz repetition rate, and a 60× oil objective was used for detection. The TCSPC decay curves were fitted with single exponential using the SymphoTime 64 software (PicoQuant, Germany), and the lifetime of each pixel was determined automatically. The lifetime image of each pixel was given a pseudo-color according to the lifetime fitted. For each individual cell, the GFP-positive membrane was selected as the region of interest (ROI), and the lifetime of each pixel within ROI was averaged. The averaged lifetime determined from multiple cells was used for the calculation of inter-protomer FRET distances.

FRET efficiencies (*E*) were calculated using the following equation. Here, *E* represents energy transfer efficiency; τ_D_ and τ_DA_ are the lifetime values of donor fluorophore in the absence and presence of the acceptor, respectively.

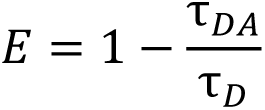

The distance between donor and acceptor was converted from FRET efficiency using the following equation. Here, *E* represents FRET efficiency, *R*_0_ is the Förster distance at which the energy transfer efficiency is 50 %, and *r* is the distance between the donor and the acceptor.

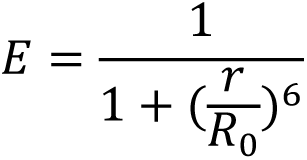

FRET distances were thus converted from FLIM lifetime measurements using GFP-mCherry Förster distance of 51 Å (***Albertazzi et al., 2009; Akrap et al., 2010; Ding et al., 2017***) and assuming κ^2^=2/3. The FRET efficiencies and FRET distances are listed in Supplementary Table 1.

### Evaluation GPR17 internalization using split-GFP labelling approach

The GFP_(10-11)_ was introduced after R291 of the wildtype or mutant GPR17 (GPR17/R291:GFP_(10-11)_), and the resulting plasmid was transfected to HEK293 cells. Two days after transfection, cells were washed with PBS and were incubated with 2 μM GFP_(1-9)_ for 20 min 37 °C, with the excess GFP_(1-9)_ washed off. The cells were cultured in DMEM with 10% fetal bovine serum for another 24 hours, in the presence or absence of 100 µM UDP-glucose. The cells were fixed by fresh prepared 4% paraformaldehyde, and mounted with an anti-fade medium containing DAPI. Imaging was taken by using the A1 confocal laser-scanning microscope, and the fluorescent intensity for a cross section of a single cell were analyzed by NIS-Elements AR software (Nikon, Tokyo, Japan).

### Evaluation of the activation of ERK1/2 by Western blotting

One day after transfection with wildtype or mutant GPR17, cells were serum starved by switching to HBS buffer (10 mM HEPES, 4 mM KCl, 140 mm NaCl, 2 mM MgSO_4_, 1 mM KH_2_PO_4_, PH 7.4) for 12 h. The cells were stimulated with 100 μM UDP-glucose for 5 min at 37 °C. The cells were then washed twice with ice-cold PBS buffer. RIPA lysis buffer (Beyotime Biotechnology, Shanghai, China) was carefully pipetted onto the well containing the adherent cultured cells. The cells were scraped off and were transferred to a microcentrifuge tube. Cell debris was removed by centrifugation at 10,000 g for 10 min at 4 °C. After quantifying protein concentration, 50 µg of the cell extracts were used for Western blotting analysis. The primary antibody included mouse anti-phospho-ERK1/2 polyclonal antibody (Cell Signaling Technology, Boston, USA, 1:1000) and rabbit anti-ERK1/2 polyclonal antibodies (Cell Signaling Technology, Boston, USA, 1:1000). The ratio of phospho-ERK1/2 over ERK1/2 indicates the activation of ERK1/2.

### Intracellular Ca^2+^-level measurement

HEK293 cells were cultured in 96 well plate, and were transfected with wildtype and mutant GPR17 plasmids as pIRES2-EGFP/GPR17-His. This plasmid can express GFP17 and GFP separately, and GFP can be used as a proxy for the expression level of GPR17. The net increase of Rhod-2 fluorescence intensity was normalized to the GFP expression level, which indicates intracellular Ca^2+^-level. 24 h after transfection, cells were incubated with 4 μM Rhod-2 (Life technologies, Eugene, USA) for 30 min at 37 °C in humidified air with 5% CO_2_. The cells were then cultured with a balanced salt solution (137 mM NaCl, 5.36 mM KCl, 1.26 mM CaCl_2_, 0.81 mM MgSO4_7_H_2_O, 0.34 mM Na_2_HPO_4_·7H_2_O,0.44 mM KH_2_PO_4_, 4.17 mM NaHCO_3_, 10 mM HEPES, and 2.02 mM glucose, pH 7.4) for another 30 min at 37 °C. The fluorescence intensity was read using appropriate wavelength settings (excitation at 550 nm, emission at 581 nm for Rhod-2; excitation at 485 nm, emission at 512 nm for GFP as expression control) on a microplate reader (Perkin-Elmer, Waltham, USA). 20, 100 and 500 µM UDP-glucose was applied sequentially, and the change of fluorescent intensity was measured continuously at 1 second intervals.

### Cell-based cAMP assay

Intracellular cAMP levels were measured using bioluminescence resonance energy transfer (BRET) assay as previously described (***Jiang et al., 2007***). Briefly, cells were co-transfected with the BRET-based cAMP biosensor CAMYEL (50 ng) and pcDNA3.1/HA-GPR17 (50 ng) or the mutant GPR17 plasmids (50 ng) per well in 96-well white plate (Thermo Scientific, Denmark). One day after transfection, cells were washed twice with PBS and incubated with 5 μM coelenterazine H (Promega, Madison, USA) for 5 min 37 °C. 2 μM Forskolin (Sigma-Aldrich, St. Louis, MO, USA) was added to stimulate adenylyl cyclase and was incubated with the cells for 5 min, in the absence or presence of 1 mM UDP-glucose. Luminescence emissions at 530 and 485 nm were measured using a Mithras LB 940 (Berthold, Bad Wildbad, Germany), and the BRET signal was presented as the 530/485 ratio.

### Statistical Analysis

The graphs and statistical data analyses were performed using GraphPad Prism 6.0 software. The results were expressed as mean ± SD for the FLIM-FRET data and mean ± SEM for other data. p<0.05 was considered statistically significant. The experiment repeat, as well as detailed statistical information including the number of experiments, p-values, definition of error bars, (t, df) and 95% CI of difference is listed in individual figure legends.

### Modeling of GPR17 protomer structure

The structure of GPR17 protomer was predicted using with threading modeling using the I-TASSER server (***Roy et al., 2010***), with the distance restraints from coevolution analysis generated with GREMLIN (***Ovchinnikov et al., 2014***). Five GPR17 models were provided by the online server with the RMSD of 0.6 Å. The GPR17 protomer structure was then subjected to MD simulations. The lipid-bilayer membrane environment was built using CHARMM-GUI Lipid Builder server(***Jo et al., 2008***). 64 DOPC molecules were inserted to either lower or upper leaflet of the bilayer and the starting thickness of the system is ∼17.5 Å. 150 mM NaCl was added to the system. The MD simulations were performed using AMBER14 (***Case et al., 2014***). The structural model was first minimized, and the system including protein, lipids, water, and ions was gradually heated. After a 5-ns equilibration, MD simulation was performed for the system for a total of 400 ns at 303 K. The temperature was controlled using Langevin thermostat while the pressure is controlled using anisotropic Berendsen barostat. The long-range electrostatics was treated with Particle Mesh Ewald method (***Essmann et al., 1995***), and the van der Waals was truncated at 10 Å. Three predicted models from I-TASSER with best scores were used to perform MD simulations. These models appear stable, as their coordinates quickly stabilized with only minor adjustments. Hence, we treated the structure model pf the protomer as a rigid body when building the dimer structure.

### Modeling GPR17 dimer structure

To assess the conformational space of fluorescent protein, mCherry (PDB code 2H5Q) and GFP (PDB code 2B3P) were inserted at specific sites, and were treated as rigid bodies. The GPR17 residues at the insertion site (the N-terminus, ECL1 after residue 128, ECL2 after residue 214, or ECL3 after residue 291), the C-terminal flexible residues of mCherry (residues 222-223), the loop residues at both N- and C-terminal ends of the inserted β-strands of GFP (residues 197-198 and 229-232), and the additionally inserted glycine residues flanking the inserted β-strands, were given full torsion angle freedom. All possible conformations of the engineered mCherry and GFP tags were randomized using in Xplor-NIH software package (***Schwieters et al., 2018***). The fluorescent proteins were found to take up a large conformational space with respect to GPR17, without clashing with GPR17 or with the lipid bilayer. The geometric center of all possible positions of the fluorophore at a particular insertion site was calculated, and the averaged position was used for the application of FLIM-FRET distance restraints (Fig. 3a).

The modeling of GPR17 dimer was performed using Xplor-NIH. The coordinates for GPR17 along with the pseudo-atoms for the averaged center-of-mass of the inserted fluorophores were duplicated. The two protomers of GPR17 were enforced with a rotational dimer symmetry. FRET distance restraints were applied to the pairs of pseudo-atoms (doubled for the *C*_2_ symmetry), and a ellipsoidal radius-of-gyration restraint (***Schwieters and Clore, 2008***) was applied for the compactness of the dimer. The calculation was performed with simulated annealing refinement similar to previously described protocol (***Ding et al., 2017***), and was repeated multiple times with different relative positions/orientations for the two protomers. The obtained GPR17 dimer structure was subjected to MD simulations to assess the stability of the coordinates, with 128 DOPC molecules were inserted to either inner or outer leaflet of the lipid bilayer. PyMOL was used to render structural figures, and to introduce point mutations for subsequent simulations (The PyMOL Molecular Graphics System, Version 2.2 Schrödinger, LLC.).

### Assessment of dimer stability with steered MD

We performed adaptive steered molecular dynamics simulations (ASMD) to obtain the energy difference between the two interacting protomers and two non-interacting separated protomers. The RMS deviation and distance between the centers-of mass was calculated using the PTRAJ module in AMBER14. The starting conformation is obtained from the last snapshot of the regular MD simulations as described above. The distance between the Cα atoms of D105 (located at the opposite side of the dimer interface) in GPR17 protomers were used for ASMD simulations, which starts from ∼44 Å to a target value of 54 Å. The velocity of ASMD is 0.5 Å/ns, and the simulations runs for a total 20 ns. The potential mean of force (PMF) was calculated, which gives the work needed to disrupt the dimer.

---

**Figure 2—figure supplement 1.**
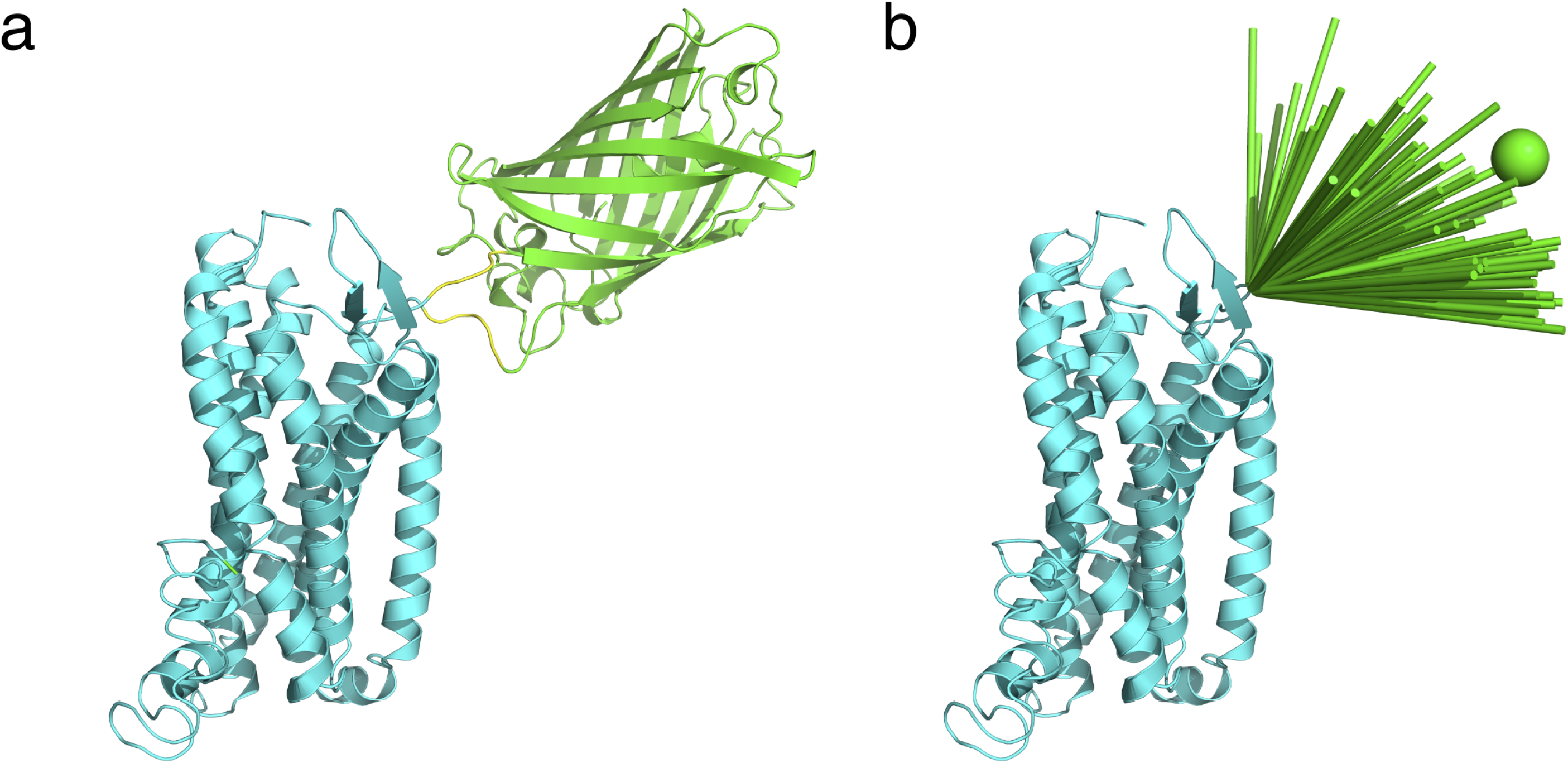
The conformational space of the fluorescent protein tagged at GPR17. (a) A representative structure of GFP inserted at ECL2 of GPR17. The GFP (PDB code 2B3P) was inserted with a split-GFP stratagem, in which the two C-terminal β-strands were engineered after GPR17 residue R214. Four glycine residues, colored yellow, flank each side the insertion. The GFP 1-9 β-stands were added to the cells to complement the already inserted strands and to regenerate fluorescence. The GFP and the GPR17 are shown as green and cyan cartoon, respectively. (b) The inserted GFP protein takes up multiple conformations with respect to GPR17. Therefore, the effect of orientation factor κ^2^ on FRET efficiency is likely small. With the conformations clashing with the membrane excluded, vectors are drawn from residue R214 to the center-of-mass of GFP in all possible conformations. The averaged position, shown as sphere, is used for subsequent modeling of dimer structure. Note that mCherry (PDB code 2H5Q) is appended at the flexible N-terminus of GPR17, which can also sample a large conformational space.

**Figure 3—figure supplement 1.**
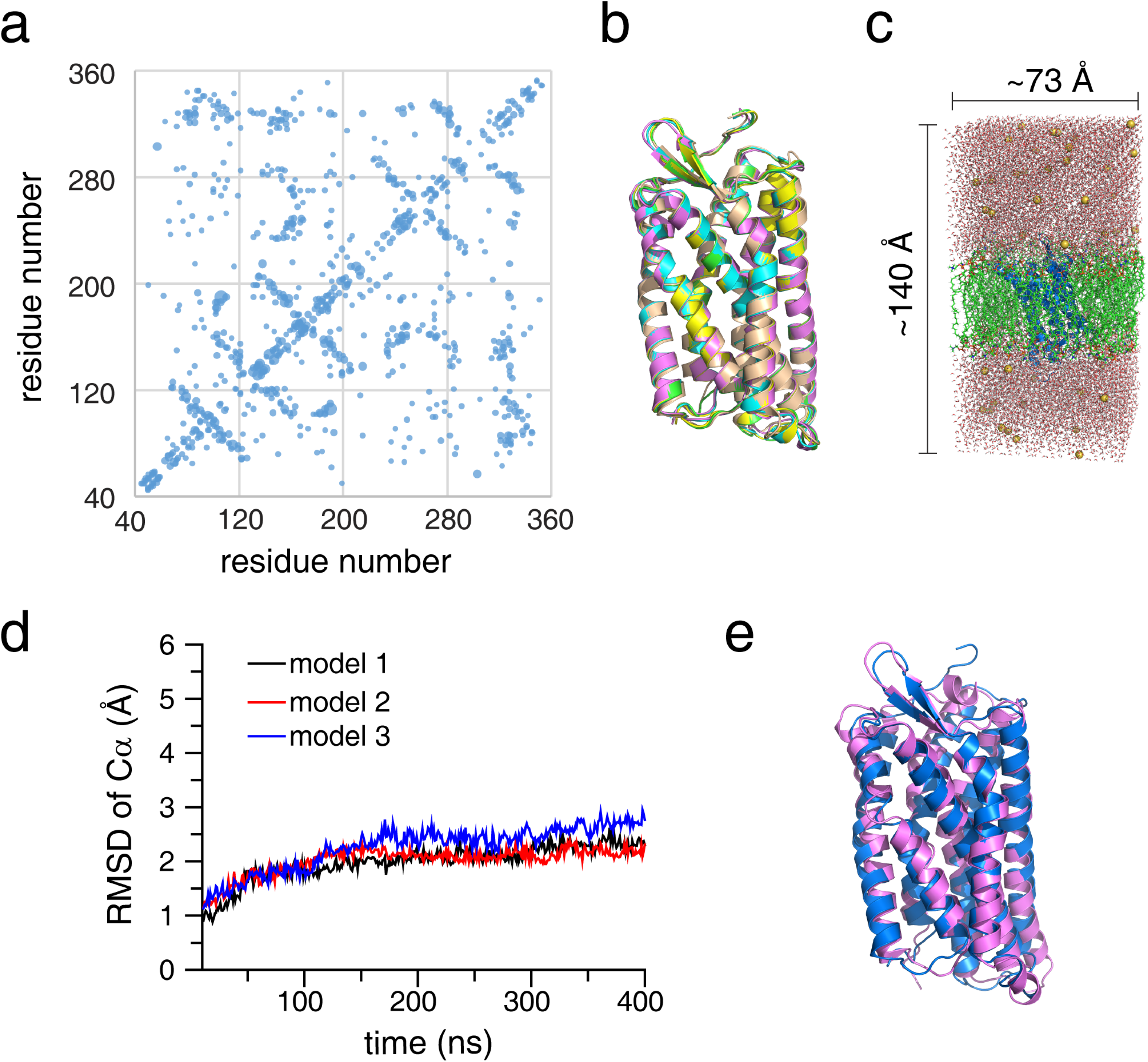
Modeling of GPR17 protomer structure. (a) Coevolution of residues of GPR17 analyzed using GREMLIN. The larger dots indicate the higher strength in covariance. Coevolution means that the two residues are likely in proximity with each other, which is used as weak distance restraints. (b) Structural models of GPR17 protomer built with I-TASSER. The three predicted models have an RMS differences of 0.61 Å. (c) MD simulation of GPR17 protomer, with sufficient amount of lipid molecules built around the protein, with water molecules placed on top and bottom of the lipid bilayer, and with periodic boundary conditions enforced. (d) RMS deviations of the Cα atoms using the three predicted models as the starting coordinates. After some small initial adjustment, all the structural models stabilize. (e) Superposition of input model and the model generated after 400 ns simulation, with the backbone RMS difference stabilizing to 2.2 Å.

**Figure 3—figure supplement 2.**
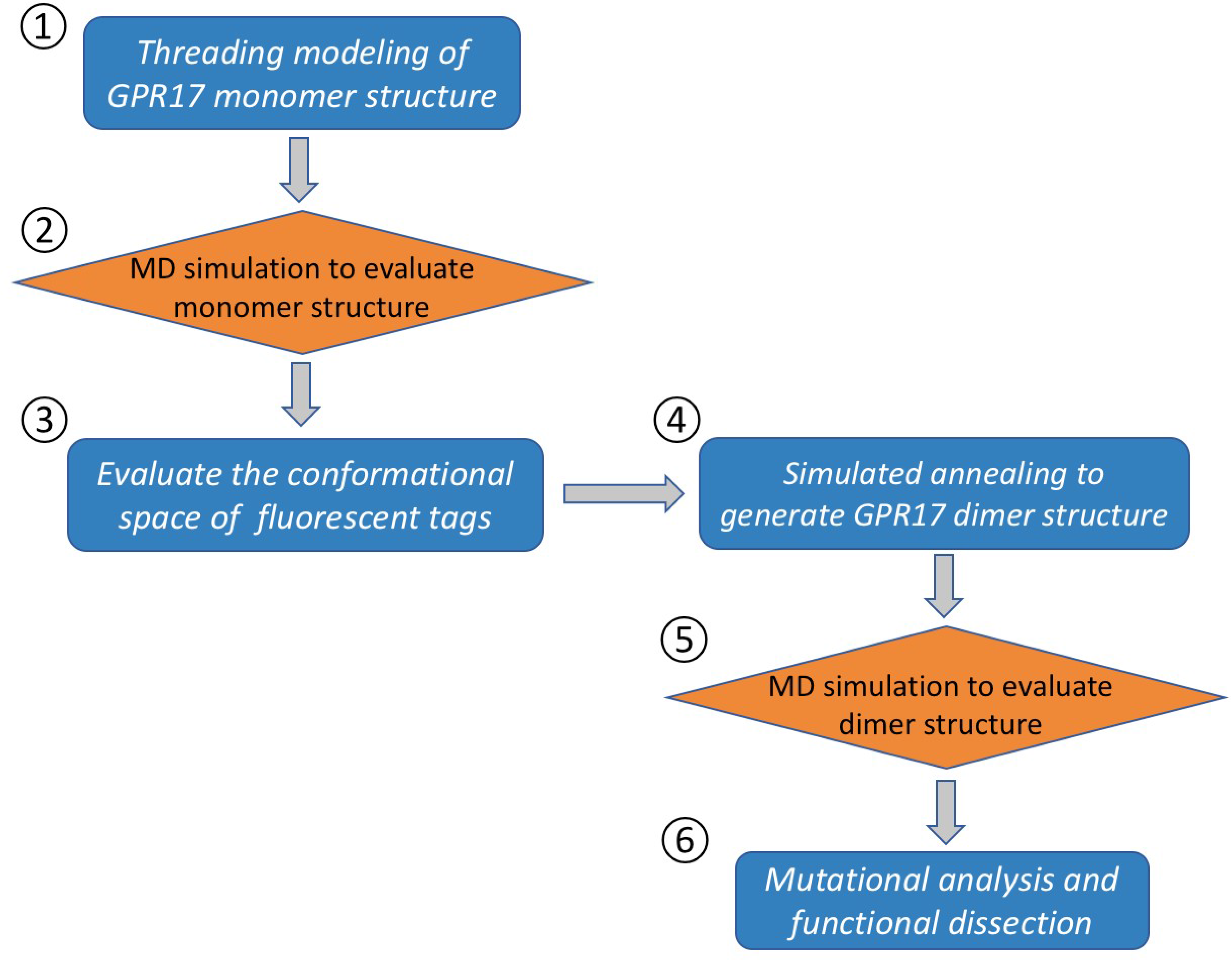
Flowchart for modeling GPR17 dimer structure, which comprises six steps. 1) Modeling of the protomer structure was performed with I-TASSER with the threading of homolog structures, and with additional input of coevolution restraints generated with GREMILIN. 2) The structural models from I-TASSER were evaluated for stability and rigidity with MD simulations using AMBER14 software package. 3) Fluorescent proteins, sfGFP or mCherry, were built into the GPR17 protomer structure, with the positions of the fluorophore with respect to GPR17 fully sampled. 4) With the averaged position of the center-of-mass of the fluorophore calculated, and with the GPR17 protomer treated as a rigid body, the GPR17 dimer structure was refined with simulated annealing in Xplor-NIH suite, so as to satisfy all FRET distance restraints between the fluorophores in the two neighboring protomers. 5) The dimer structure was further assessed with MD simulation for stability. 6) Interfacial mutations were rationally designed based on the dimer structure, and the functions of forced GPR17 monomer/dimer were evaluated.

**Figure 3—figure supplement 3.**
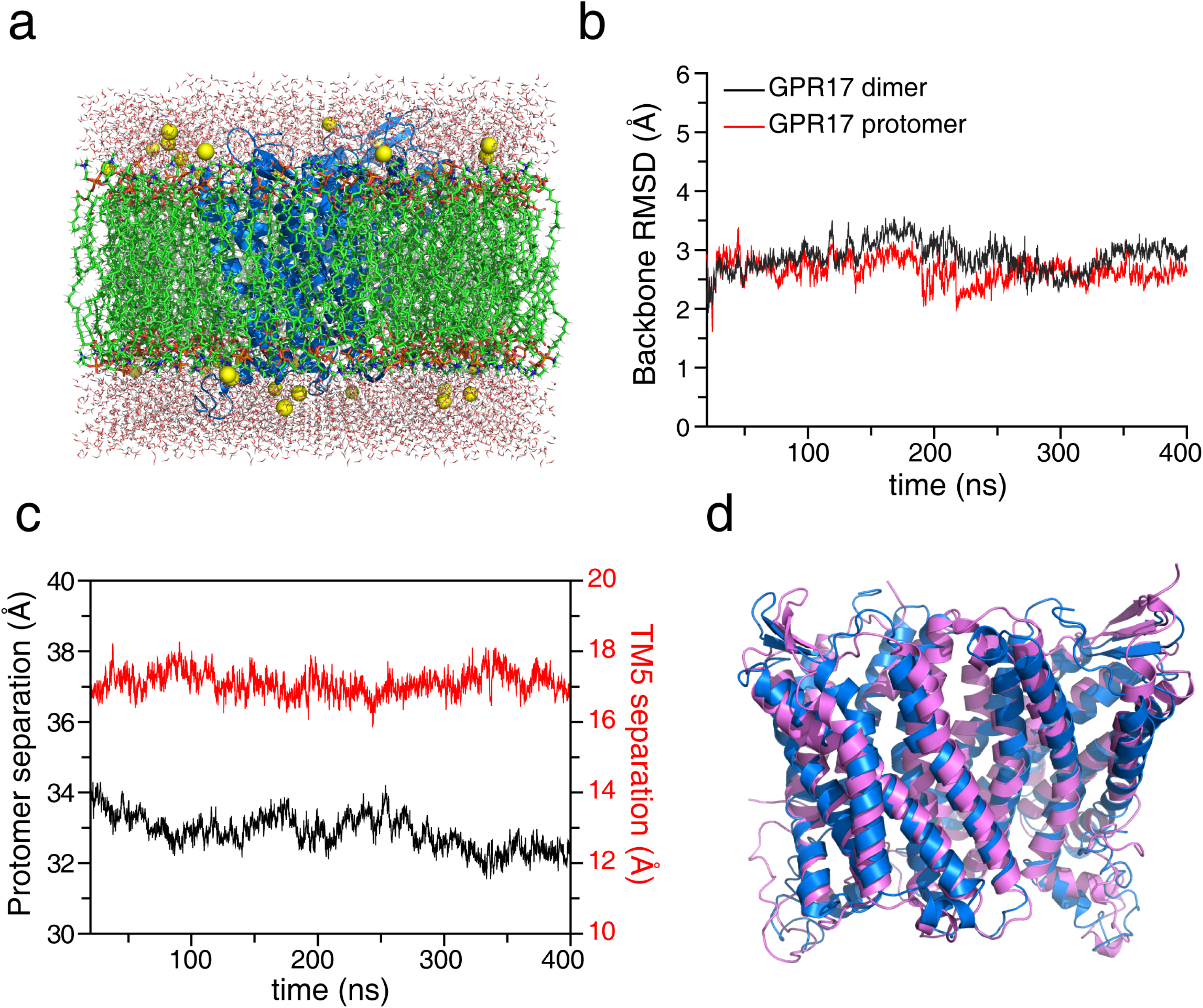
The dimer structure of GPR17 is stable in lipid bilayer. (a) GPR17 dimer structure is subjected to MD simulation. In this case, more lipid molecules are built around the protein, resulting in a larger periodic box than in the simulation for GPR17 protomer. The N-terminal extracellular tail is removed, thus to decrease the water molecules required at the top and bottom of the lipid bilayer. (b, c) After some initial adjustment, the GPR17 dimer structure stabilizes. This is characterized by similar RMS deviation of backbone Cα atoms for the dimer and for the protomer (b), and nearly constant distances between the centers-of-mass of the two protomers and of the two transmembrane helices 5 (TM5) (c). (d) Superposition of the initial input structure built with Xplor-NIH, and final MD simulated structure generated with AMBER14, affording an overall backbone RMS difference of 3.05 Å.

**Figure 3—figure supplement 4.**
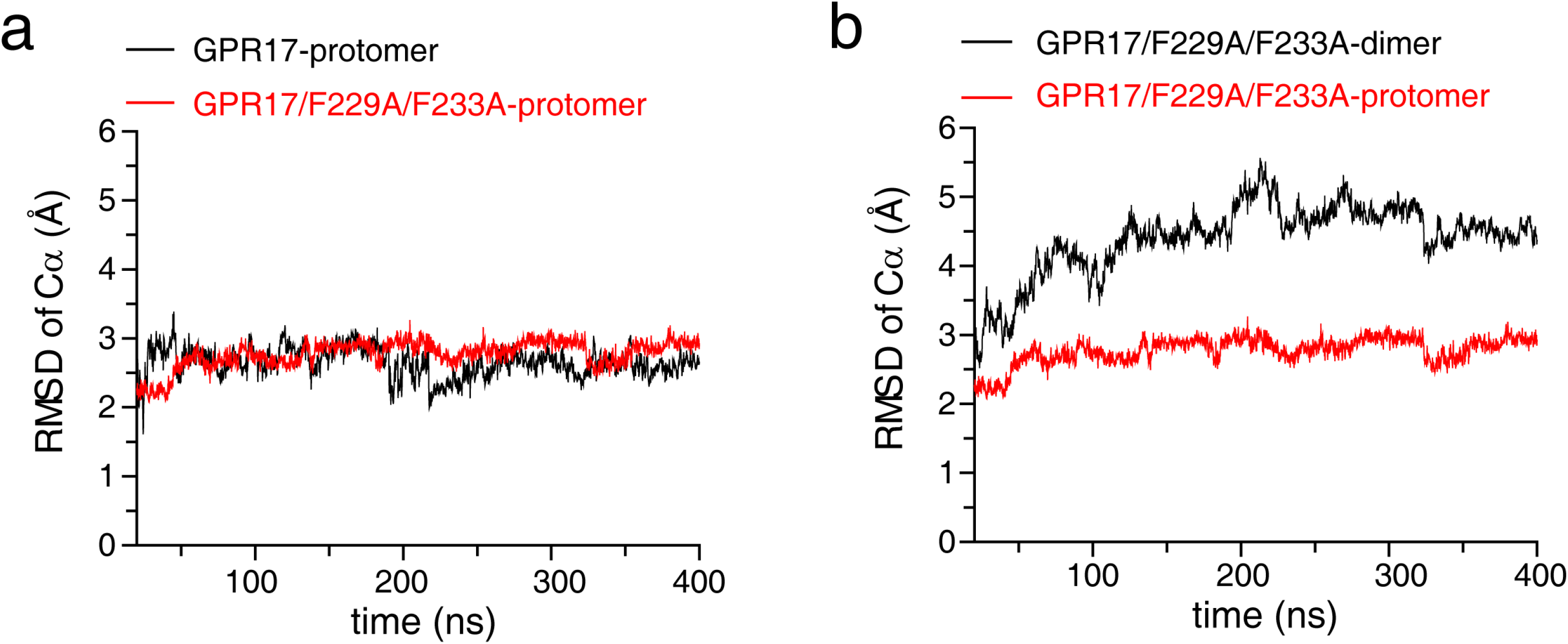
F229A/F233A mutations destabilizes GPR17 dimer. (a) With alanine mutations introduced *in silico*, the fluctuation of coordinates during MD simulations of the mutant GPR17 protomer, as characterized by the RMS deviations of backbone Cα atoms, is comparable to that of wildtype GPR17 protomer. (b) The RMS deviations of the coordinates is much larger for the mutant GPR17 dimer, and gradually increases.

**Figure 3—figure supplement 5.**
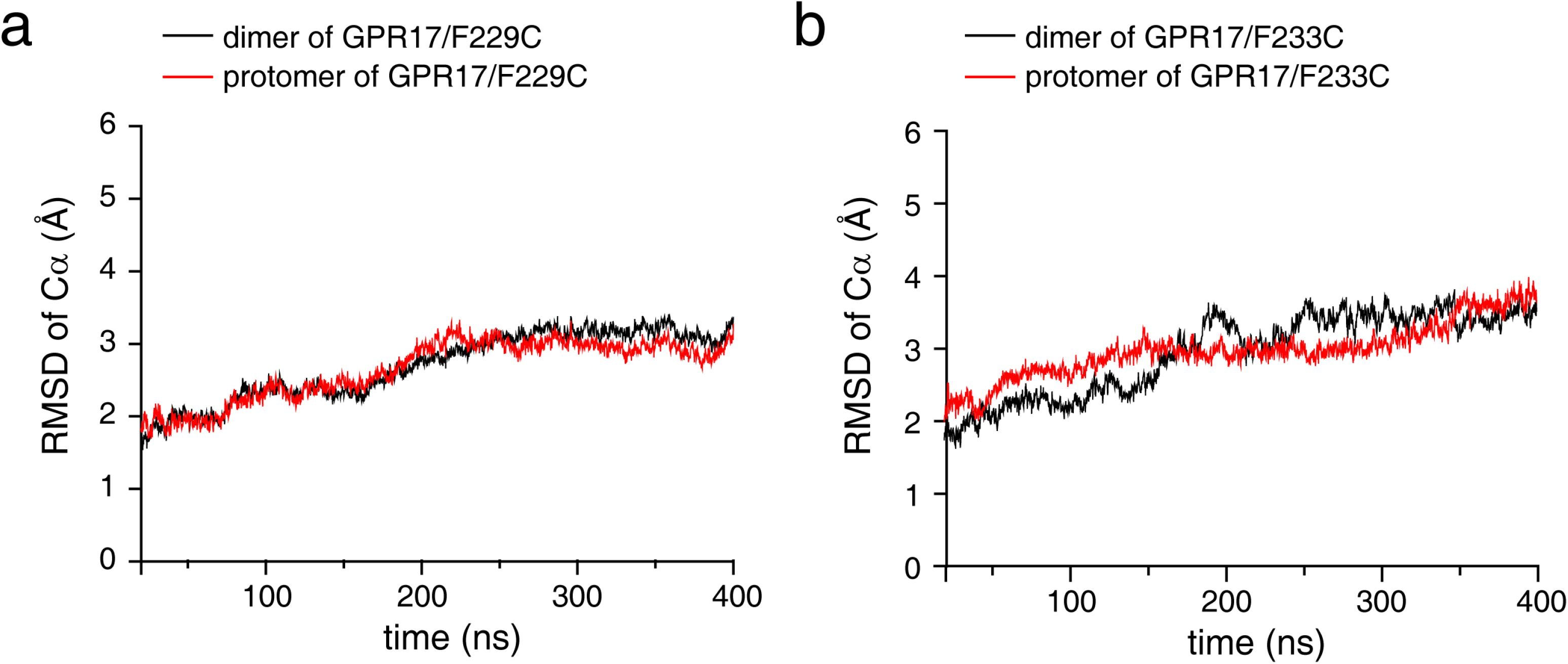
Cysteine mutation to F229 or F233 makes GPR17 dimeric. With F229 (a) or F233 (b) mutated to cysteine, the two GPR17 protomers form a disulfide-linked covalent dimer. The mutant GPR17 dimer remains stable during MD simulations, and the backbone RMS deviations are comparable with those of GPR17 protomer. This means the fluctuation of the dimer coordinates mainly arises mainly from within each protomer.

**Figure 5—figure supplement 1.**
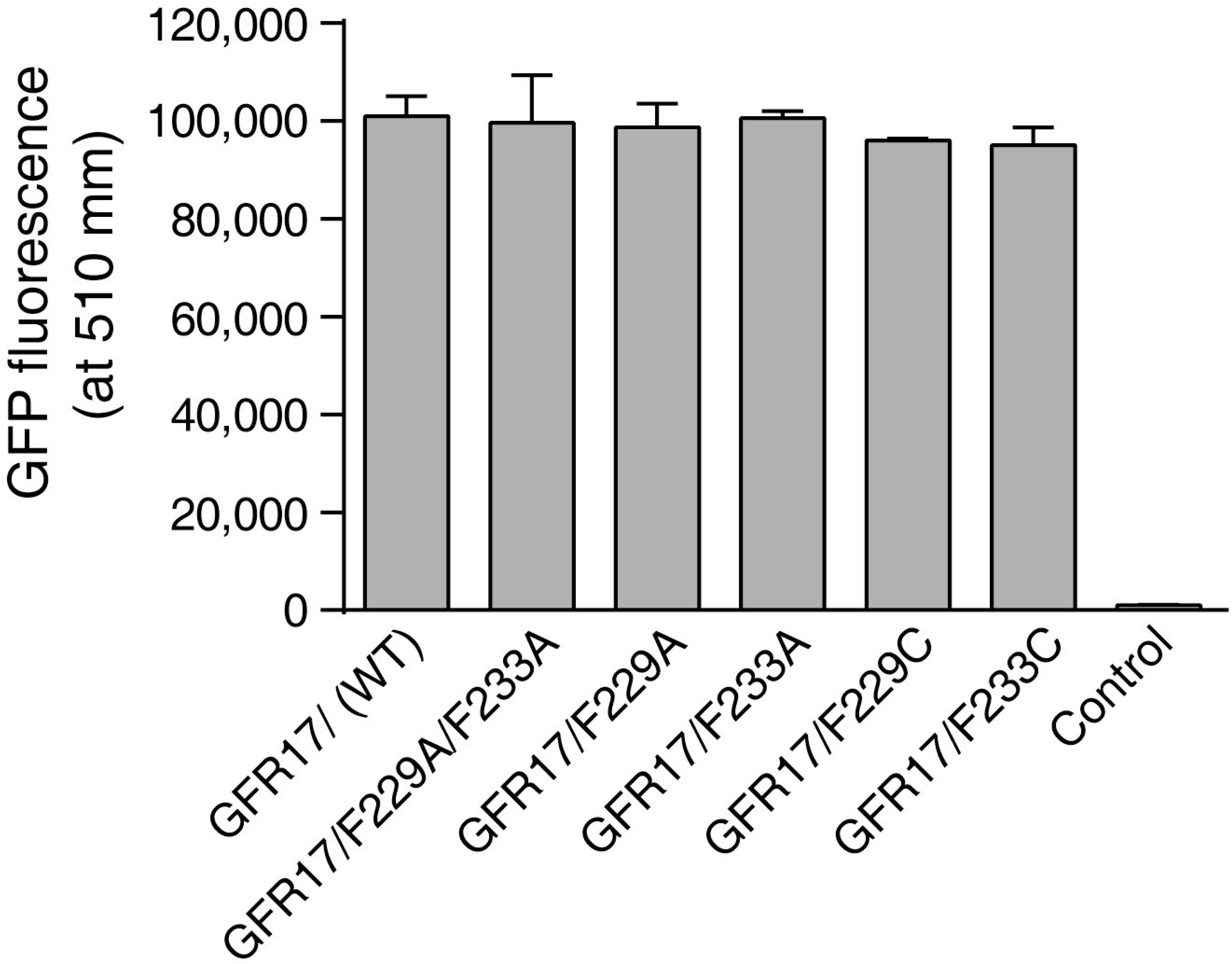
Assessment of wildtype or mutant GPR17 expression levels in HEK293 cells. Wildtype and mutant GPR17 genes were cloned into pIRES2-EGFP vector and were transfected into HEK293 cells. The pIRES2 is a bicistronic vector that allows simultaneous expression of GPR17 and GFP from the same mRNA transcript. Thus, fluorescence measurements from the GFP can be used as a proxy for the expression levels of GPR17 proteins. 24 hours after transient transfection of the plasmid, intracellular GFP fluorescence was read at 488 nm excitation and 510 nm emission. Mean ± SEM (Figure 5*—source data 2*). n= 4. One-way ANOVA followed by Tukey’s multiple comparisons test was used, with p value =0.0192 and F(6, 21)=254.0.

**Figure 6—figure supplement 1.**
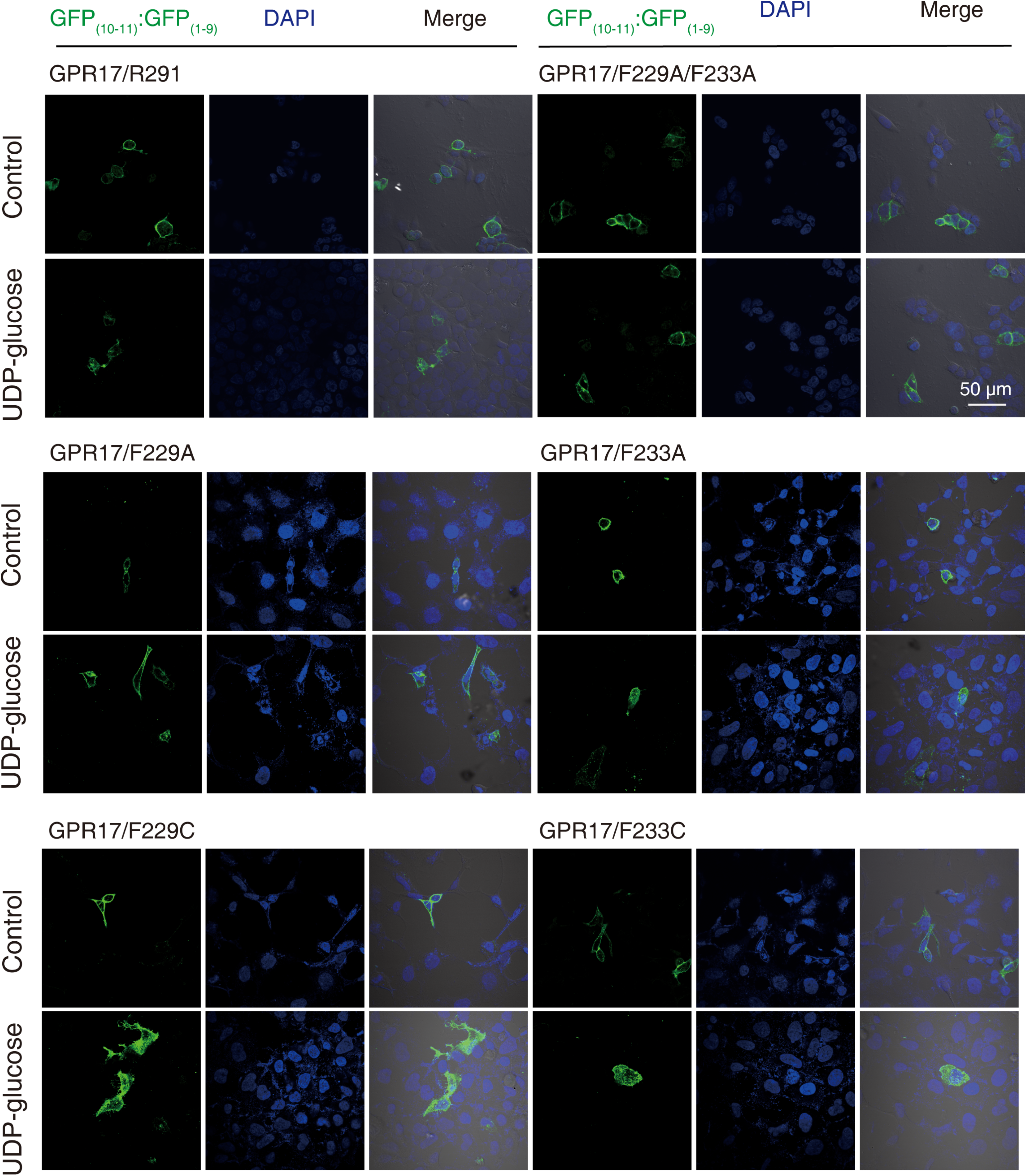
Representative images of HEK293 cells expressing GFP-tagged GPR17. The cells were transiently transfected with wildtype or mutant GPR17 gene, which carries an insertion of the C-terminal two β-strands of GFP at specific site. 24 hours after the transfection, a fragment comprising GFP β-strands 1-9 was added and was incubated with the cells for 20 minutes. This GFP fragment complements the β-strands already inserted in GPR17 expressing at cell membrane and generates GFP fluorescence at the cell surface. The cells were then treated with or without UDP-glucose for 24 hours, and the GFP fluorescence images were captured. The micrographs indicate the internalization of GFP-tagged GPR17 receptors, except for F229A and F229A/F233A mutants.

**Figure 7—figure supplement 1.**
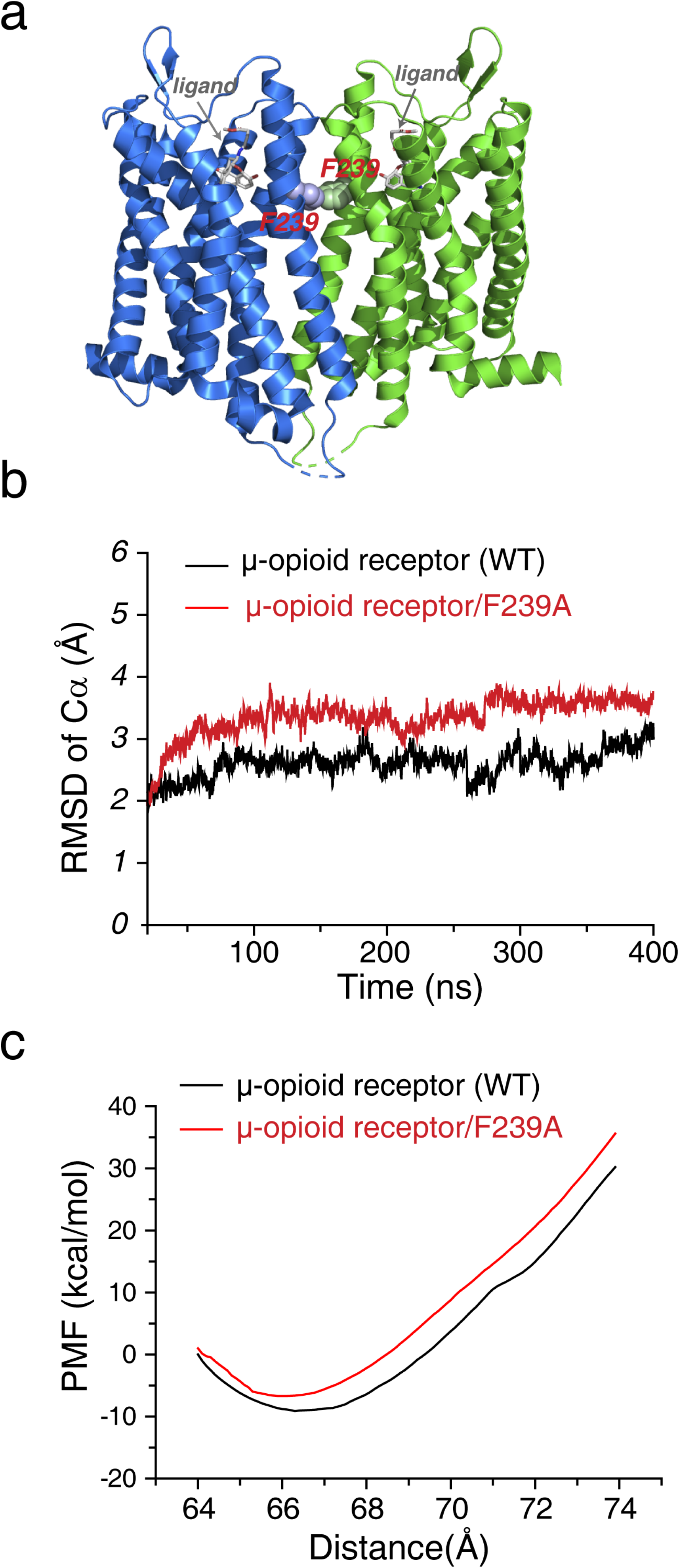
MD simulation analysis of µ-opioid receptor dimer. (a) The µ-opioid receptor has been captured as a dimer in the crystal (PDB structure 4DKL), which also utilizes TM5/6 as the dimer interface. The two protomers are arranged along the crystal *C*_2_ symmetry axis. (b) MD simulation of µ-opioid receptor dimer shows that the RMS deviations of the Cα atoms become larger upon the F239A mutaiton. (c) Steered MD simulation shows that interfacial mutaiton F239A makes µ-opioid receptor dimer less stable by ∼2 kCal/mol.

## References

Akrap N, Seidel T, Barisas BG. 2010. Forster distances for fluorescence resonant energy transfer between mCherry and other visible fluorescent proteins. Anal Biochem 402:105–106. doi:10.1016/j.ab.2010.03.026.

Albertazzi L, Arosio D, Marchetti L, Ricci F, Beltram F. 2009. Quantitative FRET analysis with the EGFP-mCherry fluorescent protein pair. Photochem Photobiol 85:287–297. doi:10.1111/j.1751-1097.2008.00435.x.

Baltoumas FA, Theodoropoulou MC, Hamodrakas SJ. 2016. Molecular dynamics simulations and structure-based network analysis reveal structural and functional aspects of G-protein coupled receptor dimer interactions. J Comput Aided Mol Des 30:489–512. doi:10.1007/s10822-016-9919-y.

Becker W. 2012. Fluorescence lifetime imaging--techniques and applications. J Microsc 247:119–136. doi:10.1111/j.1365-2818.2012.03618.x.

Berezin MY, Achilefu S. 2010. Fluorescence lifetime measurements and biological imaging. Chem Rev 110:2641–2684. doi:10.1021/cr900343z.

Bonfanti E, Gelosa P, Fumagalli M, Dimou L, Vigano F, Tremoli E, Cimino M, Sironi L, Abbracchio MP. 2017. The role of oligodendrocyte precursor cells expressing the GPR17 receptor in brain remodeling after stroke. Cell Death Dis 8:e2871. doi:10.1038/cddis.2017.256.

Buccioni M, Marucci G, Dal Ben D, Giacobbe D, Lambertucci C, Soverchia L, Thomas A, Volpini R, Cristalli G. 2011. Innovative functional cAMP assay for studying G protein-coupled receptors: application to the pharmacological characterization of GPR17. Purinergic Signal 7:463–468. doi:10.1007/s11302-011-9245-8.

Calebiro D, Rieken F, Wagner J, Sungkaworn T, Zabel U, Borzi A, Cocucci E, Zurn A, Lohse MJ. 2013. Single-molecule analysis of fluorescently labeled G-protein-coupled receptors reveals complexes with distinct dynamics and organization. Proc Natl Acad Sci U S A 110:743–748. doi:10.1073/pnas.1205798110.

Calebiro D, Sungkaworn T. 2018. Single-Molecule Imaging of GPCR Interactions. Trends Pharmacol Sci 39:109–122. doi:10.1016/j.tips.2017.10.010.

Capra V, Mauri M, Guzzi F, Busnelli M, Accomazzo MR, Gaussem P, Nisar SP, Mundell SJ, Parenti M, Rovati GE. 2017. Impaired thromboxane receptor dimerization reduces signaling efficiency: A potential mechanism for reduced platelet function in vivo. Biochem Pharmacol 124:43–56. doi:10.1016/j.bcp.2016.11.010.

Case D, Babin V, Berryman J, Betz R, Cai Q, Cerutti D, Cheatham Iii T, Darden T, Duke R, Gohlke H. 2014. Amber 14.

Ciana P, Fumagalli M, Trincavelli ML, Verderio C, Rosa P, Lecca D, Ferrario S, Parravicini C, Capra V, Gelosa P, Guerrini U, Belcredito S, Cimino M, Sironi L, Tremoli E, Rovati GE, Martini C, Abbracchio MP. 2006. The orphan receptor GPR17 identified as a new dual uracil nucleotides/cysteinyl-leukotrienes receptor. EMBO J 25:4615–4627. doi:10.1038/sj.emboj.7601341.

Daniele S, Trincavelli ML, Fumagalli M, Zappelli E, Lecca D, Bonfanti E, Campiglia P, Abbracchio MP, Martini C. 2014. Does GRK-beta arrestin machinery work as a “switch on” for GPR17-mediated activation of intracellular signaling pathways? Cell Signal 26:1310–1325. doi:10.1016/j.cellsig.2014.02.016.

Daniele S, Trincavelli ML, Gabelloni P, Lecca D, Rosa P, Abbracchio MP, Martini C. 2011. Agonist-induced desensitization/resensitization of human G protein-coupled receptor 17: a functional cross-talk between purinergic and cysteinyl-leukotriene ligands. J Pharmacol Exp Ther 338:559–567. doi:10.1124/jpet.110.178715.

Denisov IG, Sligar SG. 2017. Nanodiscs in Membrane Biochemistry and Biophysics. Chem Rev 117:4669–4713. doi:10.1021/acs.chemrev.6b00690.

Ding YH, Gong Z, Dong X, Liu K, Liu Z, Liu C, He SM, Dong MQ, Tang C. 2017. Modeling Protein Excited-state Structures from “Over-length” Chemical Cross-links. J Biol Chem 292:1187–1196. doi:10.1074/jbc.M116.761841.

Ecke D, Hanck T, Tulapurkar ME, Schafer R, Kassack M, Stricker R, Reiser G. 2008. Hetero-oligomerization of the P2Y11 receptor with the P2Y1 receptor controls the internalization and ligand selectivity of the P2Y11 receptor. Biochem J 409:107–116. doi:10.1042/BJ20070671.

Essmann U, Perera L, Berkowitz ML, Darden T, Lee H, Pedersen LG. 1995. A Smooth Particle Mesh Ewald Method. Journal of Chemical Physics 103:8577–8593. doi:Doi 10.1063/1.470117.

Faklaris O, Cottet M, Falco A, Villier B, Laget M, Zwier JM, Trinquet E, Mouillac B, Pin JP, Durroux T. 2015. Multicolor time-resolved Forster resonance energy transfer microscopy reveals the impact of GPCR oligomerization on internalization processes. FASEB J 29:2235–2246. doi:10.1096/fj.14-260059.

Garcia-Nafria J, Nehme R, Edwards PC, Tate CG. 2018. Cryo-EM structure of the serotonin 5-HT1B receptor coupled to heterotrimeric Go. Nature 558:620–623. doi:10.1038/s41586-018-0241-9.

Ge B, Lao J, Li J, Chen Y, Song Y, Huang F. 2017. Single-molecule imaging reveals dimerization/oligomerization of CXCR4 on plasma membrane closely related to its function. Sci Rep 7:16873. doi:10.1038/s41598-017-16802-7.

Gibert A, Lehmann M, Wiesner B, Schülein R (2017). The monomer/homodimer equilibrium of G protein-coupled receptors: formation in the secretory pathway and potential functional significance. G-Protein-Coupled Receptor Dimers. K. Herrick-Davis, G. Milligan and G. Di Giovanni. Cham, Switzerland, Humana Press: 359–384.

Greife A, Felekyan S, Ma Q, Gertzen CG, Spomer L, Dimura M, Peulen TO, Wohler C, Haussinger D, Gohlke H, Keitel V, Seidel CA. 2016. Structural assemblies of the di- and oligomeric G-protein coupled receptor TGR5 in live cells: an MFIS-FRET and integrative modelling study. Sci Rep 6:36792. doi:10.1038/srep36792.

Guo H, An S, Ward R, Yang Y, Liu Y, Guo XX, Hao Q, Xu TR. 2017. Methods used to study the oligomeric structure of G-protein-coupled receptors. Biosci Rep 37. doi:10.1042/BSR20160547.

Gurevich VV, Gurevich EV. 2018. GPCRs and Signal Transducers: Interaction Stoichiometry. Trends Pharmacol Sci 39:672–684. doi:10.1016/j.tips.2018.04.002.

Hauser AS, Chavali S, Masuho I, Jahn LJ, Martemyanov KA, Gloriam DE, Babu MM. 2018. Pharmacogenomics of GPCR Drug Targets. Cell 172:41–54 e19. doi:10.1016/j.cell.2017.11.033.

Hebert TE, Moffett S, Morello JP, Loisel TP, Bichet DG, Barret C, Bouvier M. 1996. A peptide derived from a beta2-adrenergic receptor transmembrane domain inhibits both receptor dimerization and activation. J Biol Chem 271:16384–16392. doi 10.1074/jbc.271.27.16384.

Hern JA, Baig AH, Mashanov GI, Birdsall B, Corrie JE, Lazareno S, Molloy JE, Birdsall NJ. 2010. Formation and dissociation of M1 muscarinic receptor dimers seen by total internal reflection fluorescence imaging of single molecules. Proc Natl Acad Sci U S A 107:2693–2698. doi:10.1073/pnas.0907915107.

Isberg V, de Graaf C, Bortolato A, Cherezov V, Katritch V, Marshall FH, Mordalski S, Pin JP, Stevens RC, Vriend G, Gloriam DE. 2015. Generic GPCR residue numbers - aligning topology maps while minding the gaps. Trends Pharmacol Sci 36:22–31. doi:10.1016/j.tips.2014.11.001.

Ito T, Oshita S, Nakabayashi T, Sun F, Kinjo M, Ohta N. 2009. Fluorescence lifetime images of green fluorescent protein in HeLa cells during TNF-alpha induced apoptosis. Photochem Photobiol Sci 8:763–767. doi:10.1039/b902341k.

Jastrzebska B, Chen Y, Orban T, Jin H, Hofmann L, Palczewski K. 2015. Disruption of Rhodopsin Dimerization with Synthetic Peptides Targeting an Interaction Interface. J Biol Chem 290:25728–25744. doi:10.1074/jbc.M115.662684.

Jiang LI, Collins J, Davis R, Lin KM, DeCamp D, Roach T, Hsueh R, Rebres RA, Ross EM, Taussig R, Fraser I, Sternweis PC. 2007. Use of a cAMP BRET sensor to characterize a novel regulation of cAMP by the sphingosine 1-phosphate/G13 pathway. J Biol Chem 282:10576–10584. doi:10.1074/jbc.M609695200.

Jiang WX, Dong X, Jiang J, Yang YH, Yang J, Lu YB, Fang SH, Wei EQ, Tang C, Zhang WP. 2016. Specific cell surface labeling of GPCRs using split GFP. Sci Rep 6:20568. doi:10.1038/srep20568.

Jo S, Kim T, Iyer VG, Im W. 2008. CHARMM-GUI: a web-based graphical user interface for CHARMM. J Comput Chem 29:1859–1865. doi:10.1002/jcc.20945.

Kasai RS, Kusumi A. 2014. Single-molecule imaging revealed dynamic GPCR dimerization. Curr Opin Cell Biol 27:78–86. doi:10.1016/j.ceb.2013.11.008.

Koehl A, Hu H, Feng D, Sun B, Zhang Y, Robertson MJ, Chu M, Kobilka TS, Laeremans T, Steyaert J, Tarrasch J, Dutta S, Fonseca R, Weis WI, Mathiesen JM, Skiniotis G, Kobilka BK. 2019. Structural insights into the activation of metabotropic glutamate receptors. Nature 566:79–84. doi:10.1038/s41586-019-0881-4.

Lambert NA, Javitch JA. 2014. CrossTalk opposing view: Weighing the evidence for class A GPCR dimers, the jury is still out. J Physiol 592:2443–2445. doi:10.1113/jphysiol.2014.272997.

Lohse MJ. 2010. Dimerization in GPCR mobility and signaling. Curr Opin Pharmacol 10:53–58. doi:10.1016/j.coph.2009.10.007.

Longo PA, Kavran JM, Kim MS, Leahy DJ. 2013. Transient mammalian cell transfection with polyethylenimine (PEI). Methods Enzymol 529:227–240. doi:10.1016/B978-0-12-418687-3.00018-5.

Lu C, Dong L, Zhou H, Li Q, Huang G, Bai SJ, Liao L. 2018. G-Protein-Coupled Receptor Gpr17 Regulates Oligodendrocyte Differentiation in Response to Lysolecithin-Induced Demyelination. Sci Rep 8:4502. doi:10.1038/s41598-018-22452-0.

Manglik A, Kruse AC, Kobilka TS, Thian FS, Mathiesen JM, Sunahara RK, Pardo L, Weis WI, Kobilka BK, Granier S. 2012. Crystal structure of the micro-opioid receptor bound to a morphinan antagonist. Nature 485:321–326. doi:10.1038/nature10954.

Marucci G, Dal Ben D, Lambertucci C, Santinelli C, Spinaci A, Thomas A, Volpini R, Buccioni M. 2016. The G Protein-Coupled Receptor GPR17: Overview and Update. ChemMedChem 11:2567–2574. doi:10.1002/cmdc.201600453.

Maurel D, Comps-Agrar L, Brock C, Rives ML, Bourrier E, Ayoub MA, Bazin H, Tinel N, Durroux T, Prezeau L, Trinquet E, Pin JP. 2008. Cell-surface protein-protein interaction analysis with time-resolved FRET and snap-tag technologies: application to GPCR oligomerization. Nat Methods 5:561–567. doi:10.1038/nmeth.1213.

McMillin SM, Heusel M, Liu T, Costanzi S, Wess J. 2011. Structural basis of M3 muscarinic receptor dimer/oligomer formation. J Biol Chem 286:28584–28598. doi:10.1074/jbc.M111.259788.

Milligan G, Ward RJ, Marsango S. 2019. GPCR homo-oligomerization. Curr Opin Cell Biol 57:40–47. doi:10.1016/j.ceb.2018.10.007.

Muto T, Tsuchiya D, Morikawa K, Jingami H. 2007. Structures of the extracellular regions of the group II/III metabotropic glutamate receptors. Proc Natl Acad Sci U S A 104:3759–3764. doi:10.1073/pnas.0611577104.

Ovchinnikov S, Kamisetty H, Baker D. 2014. Robust and accurate prediction of residue-residue interactions across protein interfaces using evolutionary information. Elife 3:e02030. doi:10.7554/eLife.02030.

Pandy-Szekeres G, Munk C, Tsonkov TM, Mordalski S, Harpsoe K, Hauser AS, Bojarski AJ, Gloriam DE. 2018. GPCRdb in 2018: adding GPCR structure models and ligands. Nucleic Acids Res 46:D440–D446. doi:10.1093/nar/gkx1109.

Romei MG, Boxer SG. 2019. Split Green Fluorescent Proteins: Scope, Limitations, and Outlook. Annu Rev Biophys. doi:10.1146/annurev-biophys-051013-022846.

Roy A, Kucukural A, Zhang Y. 2010. I-TASSER: a unified platform for automated protein structure and function prediction. Nat Protoc 5:725–738. doi:10.1038/nprot.2010.5.

Schwieters CD, Bermejo GA, Clore GM. 2018. Xplor-NIH for molecular structure determination from NMR and other data sources. Protein Sci 27:26–40. doi:10.1002/pro.3248.

Schwieters CD, Clore GM. 2008. A pseudopotential for improving the packing of ellipsoidal protein structures determined from NMR data. J Phys Chem B 112:6070–6073. doi:10.1021/jp076244o.

Shen Z, Yang X, Chen Y, Shi L. 2018. CAPA periviscerokinin-mediated activation of MAPK/ERK signaling through Gq-PLC-PKC-dependent cascade and reciprocal ERK activation-dependent internalized kinetics of Bom-CAPA-PVK receptor 2. Insect Biochem Mol Biol 98:1–15. doi:10.1016/j.ibmb.2018.04.007.

Smith TH, Li JG, Dores MR, Trejo J. 2017. Protease-activated receptor-4 and purinergic receptor P2Y12 dimerize, co-internalize, and activate Akt signaling via endosomal recruitment of beta-arrestin. J Biol Chem 292:13867–13878. doi:10.1074/jbc.M117.782359.

Stenkamp RE. 2018. Identifying G protein-coupled receptor dimers from crystal packings. Acta Crystallogr D Struct Biol 74:655–670. doi:10.1107/S2059798318008136.

Suhling K, Siegel J, Phillips D, French PM, Leveque-Fort S, Webb SE, Davis DM. 2002. Imaging the environment of green fluorescent protein. Biophys J 83:3589–3595. doi:10.1016/S0006-3495(02)75359-9.

Sun Y, Rombola C, Jyothikumar V, Periasamy A. 2013. Forster resonance energy transfer microscopy and spectroscopy for localizing protein-protein interactions in living cells. Cytometry A 83:780–793. doi:10.1002/cyto.a.22321.

Ward RJ, Xu TR, Milligan G. 2013. GPCR oligomerization and receptor trafficking. Methods Enzymol 521:69–90. doi:10.1016/B978-0-12-391862-8.00004-1.

Xiang J, Chun E, Liu C, Jing L, Al-Sahouri Z, Zhu L, Liu W. 2016. Successful Strategies to Determine High-Resolution Structures of GPCRs. Trends Pharmacol Sci 37:1055–1069. doi:10.1016/j.tips.2016.09.009.

Xue L, Rovira X, Scholler P, Zhao H, Liu J, Pin JP, Rondard P. 2015. Major ligand-induced rearrangement of the heptahelical domain interface in a GPCR dimer. Nat Chem Biol 11:134–140. doi:10.1038/nchembio.1711.

